# RoDPAA: A Robust Dissimilarity-Based Principal Angle Analysis Framework for Multi-View Microbiome Data Integration

**DOI:** 10.64898/2026.10.29.762935

**Authors:** Yanyan Xu, Yue Wang

## Abstract

Multiview microbiome studies profile the same participants across body sites or molecular layers, but integration is hampered by sparse, compositional data, modest sample sizes and methods that recognize only exact sharing. We present RoDPAA (Robust Dissimilarity-based Principal Angle Analysis), a framework that represents each view through any data-appropriate dissimilarity, such as Bray–Curtis, without log-ratio transformation or likelihood assumptions. RoDPAA aligns view-specific signal subspaces through principal angles, captures complete and partial sharing, and tests by permutation how many components are shared, using only eigendecomposition and little tuning. In Gaussian simulations, RoDPAA matched JIVE and MOFA given true ranks, ran faster and alone recovered the shared dimension in every replicate. In two inflammatory bowel disease (IBD) cohorts with sparse, zero-inflated data, Bray–Curtis-based RoDPAA recovered shared structure that was stable to prevalence filtering, whereas log-ratio-based alternatives were not. Its leading microbiome–metabolome factors, learned without disease labels, were the only ones associated with IBD.

## 1 Introduction

The gut microbiome is an important component of human health and disease [1, 2] and interacts with host physiology and metabolism across anatomical sites and molecular layers [3]. Multiview microbiome studies therefore collect complementary measurements from the same individuals, such as microbiome profiles from different body sites [1] or microbial profiles paired with metabolomic or other host molecular data [3, 4]. A central goal is to identify variation shared across these views, which may represent disease-associated biology more coherently than any single view alone. Ideally, this shared variation should be learned without reference to clinical outcomes, so that any later association with disease provides independent evidence of its biological relevance.

Unsupervised multivariate representation learning provides a natural starting point for this task. In single-view microbiome studies, principal coordinate analysis (PCoA) of beta-diversity dissimilarities such as Bray–Curtis is the standard way to summarize between-sample variation without prespecifying an outcome [5–7]. Multiview methods extend this idea to variation shared across views, but each class of method fits micro-biome data poorly. Linear methods such as canonical correlation analysis (CCA) [8] and JIVE/AJIVE [9, 10] operate in Euclidean feature space, so microbial abundances must first be transformed, for example by log-ratio transformations [11], which require zero replacement and often prevalence filtering [12]. Yet there is no consensus on how zeros should be handled, which low-prevalence taxa should be removed, or which transformation should be applied [13–15], and these choices are known to materially affect conclusions in single-view microbiome analyses [16, 17]. In multiview integration, these choices may determine not only which shared structure is recovered but whether any is detected at all. Probabilistic latent-variable models, such as iCluster and MOFA [18, 19], instead require a likelihood to be specified for each view. Standard likelihoods do not describe sparse, compositional relative abundances, so fitting the raw data risks misspecification, whereas transforming them first reintroduces the preprocessing choices above. Deep-learning methods, such as the autoencoder-based maui [20], avoid distributional assumptions but have many tuning parameters whose reliable selection requires large training samples. Such sample sizes are rarely available in multiview microbiome studies, which commonly combine thousands of microbial or molecular features with only tens to a few hundred participants [3, 4].

A separate challenge concerns the definition of shared structure itself. Many multi-view methods define sharing through an exactly common latent subspace. JIVE models joint variation as a low-rank component with identical subject scores across views [9], and AJIVE uses principal angles only to decide which directions are consistent with such a subspace [10]. Directions that are related but not identical across views are therefore assigned to view-specific variation, even when they show substantial cross-view agreement. Yet because different views capture complementary aspects of the same biology, and sequencing noise perturbs the structure estimated within each view, genuinely shared signals are more likely to be partially than exactly aligned. Partial sharing therefore needs to be tested statistically, rather than either forced into a common subspace or discarded.

A related challenge is determining how many shared components the data support. Methods such as CCA and MOFA typically fix this number in advance or choose it heuristically [19]. JIVE and AJIVE estimate the joint rank more formally [9, 10], but only for an exactly common subspace. Shared variation carried by partially aligned directions is therefore not counted, and the shared dimension may be underestimated.

Together, these limitations call for an unsupervised integration framework that respects the natural geometry of each view, requires few modeling choices, defines sharing flexibly, and tests, rather than assumes, how many shared components are present.

Here we introduce RoDPAA (Robust Dissimilarity-based Principal Angle Analysis), a dissimilarity-based principal-angle framework that meets these requirements (Fig. 1). Each view is first represented by a user-specified dissimilarity that is appropriate for that data type, such as Bray–Curtis for microbiome profiles, from which RoDPAA extracts a low-dimensional sample-level signal subspace. It then pairs the most closely aligned directions of the two signal subspaces through principal angles and quantifies the strength of sharing by the cosine of each angle. A permutation test that disrupts subject correspondence across views then determines which alignments, whether complete or partial, exceed chance, so the shared dimension is inferred from the data rather than specified in advance. Because eigendecomposition is applied only to sample-by-sample similarity matrices, whose size depends on the number of participants rather than the number of features, RoDPAA requires little tuning and is highly efficient in the modest-sample settings typical of multiview microbiome studies. By accepting user-specified pairwise dissimilarities as its input, RoDPAA naturally extends the ecological distance toolkit that underpins single-view PCoA analysis—Bray–Curtis, UniFrac, and related metrics—into the multiview setting, without requiring transformation to Euclidean feature space.

**Fig. 1.**
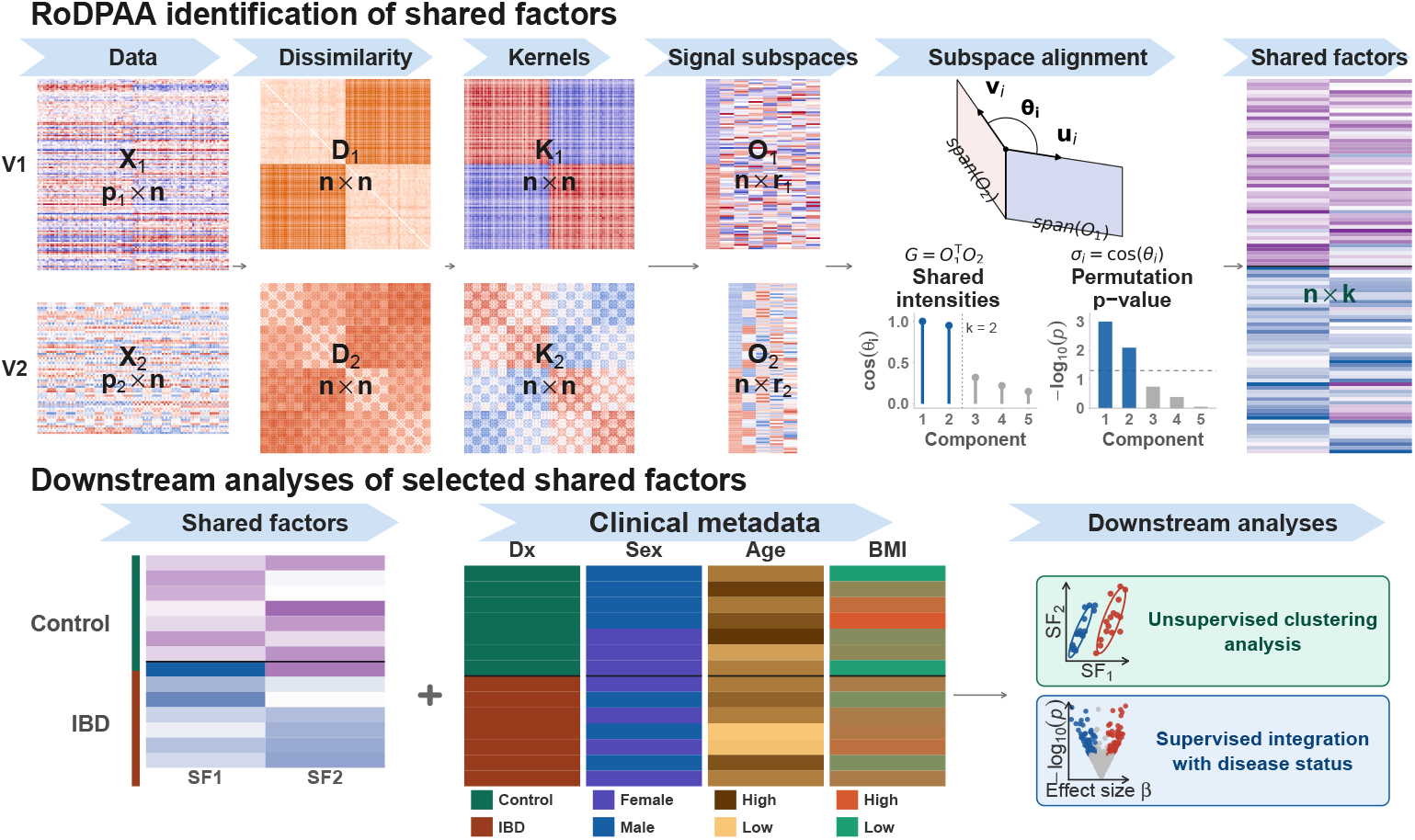
Overview of the RoDPAA workflow for multi-view biomedical data integration. RoDPAA is designed for settings in which two high-dimensional data views are measured on the same set of *n* subjects. The two input data matrices are denoted by *X*_1_ and *X*_2_, shown as *p*_1_ *× n* and *p*_2_ *× n* feature-by-subject matrices, where *p*_1_ and *p*_2_ are the numbers of features in the two views. For each view, pairwise sam-ple distances are computed to summarize subject-to-subject dissimilarity within that modality, producing dissimilarity matrices *D*_1_ and *D*_2_. These are transformed into centered kernel matrices *K*_1_ and *K*_2_ via double centering, providing sample-level similarity representations that accommodate modality-specific distance choices. Eigendecompo-sition is applied separately to each kernel matrix and the leading eigenvectors are retained as view-specific signal subspaces 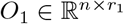 and 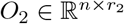, where *r*_1_ and *r*_2_ are the selected signal ranks. To identify shared structure, RoDPAA computes the principal angles (*θ*_*i*_) between the two subspaces via the cross-Gram matrix 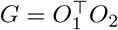. The singular values of *G* are the shared intensities, which quantify subspace alignment as *σ*_*i*_ = cos(*θ*_*i*_). A permutation-based test determines which aligned components exceed chance alignment, yielding an estimated joint rank *k*. The *k* significant aligned components are averaged across views to produce shared factors *SF*_*J*_ ℝ^*n×k*^, a low-dimensional representation of cross-view shared variation. For downstream analysis, the estimated shared factor matrix is combined with clinical metadata (disease status, sex, age, BMI): (i) unsupervised *k*-means clustering evaluations and (ii) supervised analyses that incorporate disease status and other clinical covariates to assess the biological and clinical relevance of the shared variation.

In Gaussian simulations under Euclidean geometry with known ranks, RoDPAA recovered shared group structure as accurately as JIVE and MOFA, but 5–130 times faster. When the ranks had to be estimated, RoDPAA was the only method to recover the shared dimension in every replicate under complete and partial sharing. In two IBD cohorts, Bray–Curtis-based RoDPAA recovered shared structure that was stable to prevalence filtering. By contrast, RoDPAA with Euclidean distance after centered log-ratio (CLR) transformation detected no shared microbiome–metabolome directions before filtering and 67 after, and MOFA fitted to CLR-transformed data was similarly sensitive. Although RoDPAA was fitted without disease labels, its two most strongly aligned microbiome–metabolome factors were the only ones associated with IBD.

## 2 Results

### 2.1 Overview of the RoDPAA Framework

As illustrated in Fig. 1, RoDPAA first converts the dissimilarity matrix for each view into a centered sample–sample similarity matrix and estimates its low-dimensional signal subspace from the leading eigenstructure. Principal-angle analysis of the estimated subspaces then produces paired cross-view directions and quantifies their alignment through the corresponding singular values, termed shared intensities. A permutation test that breaks subject correspondence across views while preserving the within-view signal structure then identifies the directions whose alignment exceeds that expected by chance; their number is the estimated shared dimension. The significant direction pairs are averaged across views to give participant-level shared factors. Under Euclidean geometry and exact cross-view alignment, RoDPAA reduces to the classical shared-subspace setting of JIVE and AJIVE (Methods; Supplementary Note 1); outside this setting, it extends the same framework to non-Euclidean data and partially aligned shared structure.

### 2.2 RoDPAA accurately recovers shared group structure across noise and view-specific variation

We first evaluated RoDPAA in simulations in which the shared structure was known by construction. Data were generated from a Gaussian latent-factor model, and RoDPAA used Euclidean distance, the geometry assumed by the competing methods, so that differences reflect how shared structure is estimated rather than the choice of dissimilarity (Methods). Simulation I evaluated recovery of a two-dimensional group-associated shared representation under varying levels of measurement noise and view-specific variation (Fig. 2). We compared RoDPAA with JIVE, CCA, MOFA, PCA applied separately to each view, and PCA applied to the concatenated data matrix. The target two-dimensional representation was prespecified, and JIVE and MOFA were given the true ranks (Methods), so that the comparison focused on recovery of the shared group structure rather than selection of its dimension. All methods were fitted without access to the group labels, which were used only afterward to quantify separation of the recovered participant representations using the Calinski–Harabasz (CH) index.

**Fig. 2.**
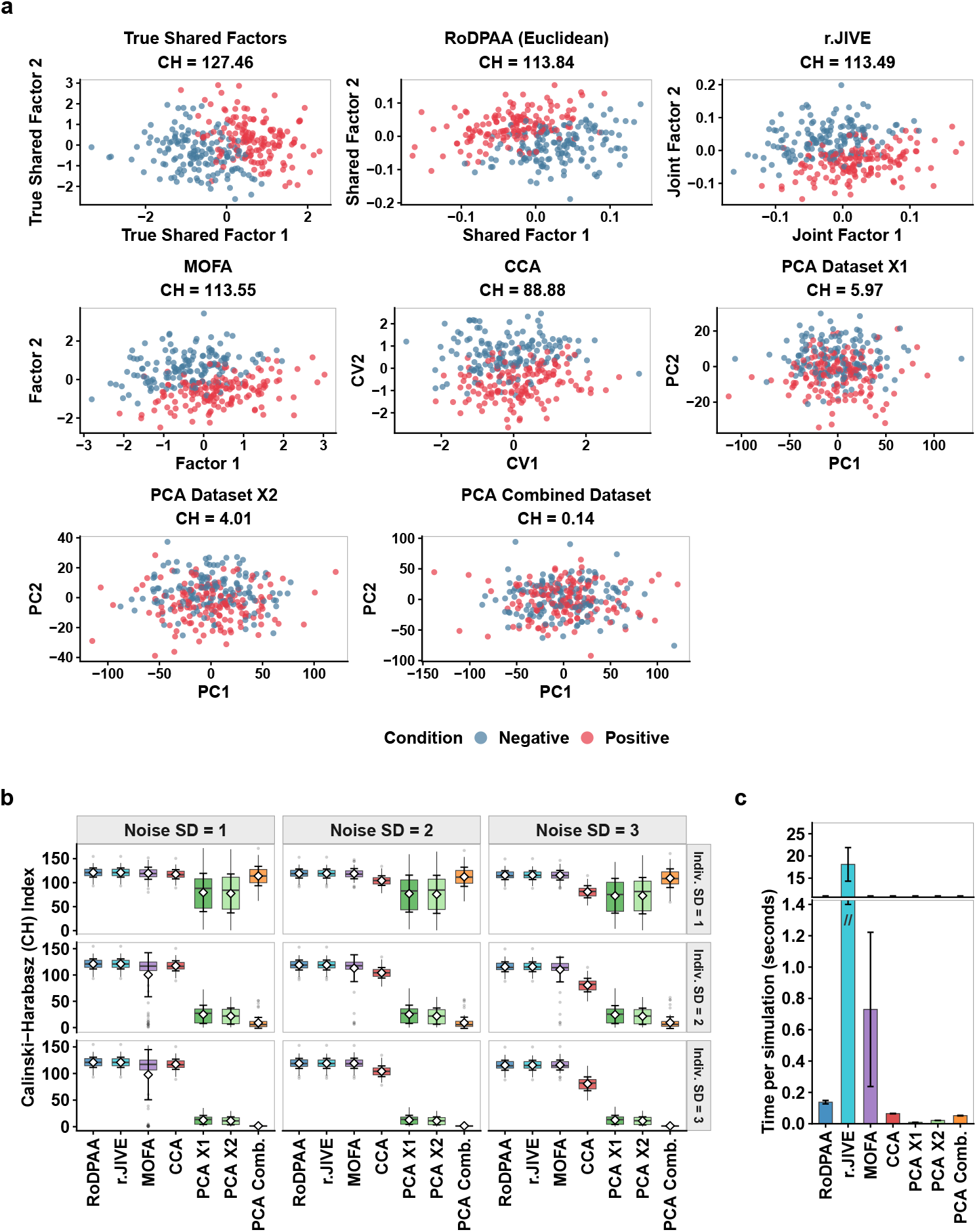
Simulation I: Recovery of the shared latent representation. **a**, Representative simulation under *σ*_noise_ = 3 and *σ*_indiv_ = 3. The first two recovered shared factors are shown for each method. Points are colored according to the true binary disease groups (red = positive, blue = negative). The Calinski–Harabasz (CH) index is reported for each recovered two-dimensional representation. JIVE was fitted with joint rank two and individual rank one per view, and MOFA with four factors, of which the first two shared factors are shown. **b**, Distribution of CH indices over 100 independent simulation replicates for each combination of *σ*_noise_ *∈ {*1, 2, 3*}* (columns) and *σ*_indiv_ ∈ {1, 2, 3} (rows). Higher CH values indicate better recovery of the disease-discriminating shared latent structure. Boxes show the median and interquartile range (whiskers, 1.5 times the interquartile range; points, outliers), and white diamonds with error bars show the mean *±* one standard deviation. Column and row strips give *σ*_noise_ (Noise SD) and *σ*_indiv_ (Indiv. SD). **c**, Mean computation time per simulation replicate averaged over all 900 simulated datasets. Error bars denote one standard deviation. The vertical axis is broken (0–1.4 s and 11–26 s) to accommodate the substantially longer runtime of r.jive. Simulation settings were *n* = 300, *p*_1_ = 100, *p*_2_ = 150, *k* = 2, *q*_1_ = *q*_2_ = 1, and *ϕ* = 0.3.

Figure 2a shows a representative dataset with *σ*_noise_ = *σ*_indiv_ = 3. The true group-associated latent factors had a CH index of 127.46. RoDPAA, JIVE and MOFA recovered clear and nearly identical group separation, with CH indices of 113.84, 113.49 and 113.55, respectively. This agreement is expected: under Euclidean geometry and with the same signal and joint ranks, RoDPAA and JIVE target the same population signal subspaces (Methods; Supplementary Note 1), and MOFA with a Gaussian likelihood can be viewed as a generative model for the JIVE decomposition [19]. CCA produced moderate group separation (CH =88.88), whereas PCA, applied to each view separately or to the concatenated data, produced little (CH 0.14–5.97).

Across 100 replicates for each of nine combinations of *σ*_noise_ and *σ*_indiv_ (Fig. 2b), concatenated PCA performed well when *σ*_indiv_ = 1 (mean CH 109.3–113.8) but declined sharply as *σ*_indiv_ increased (9.0–9.3 at *σ*_indiv_ = 2 and 1.5–1.6 at *σ*_indiv_ = 3); PCA applied to each view separately declined similarly. Because PCA targets directions of maximal total variance, its leading components become increasingly dominated by the stronger view-specific variation rather than the weaker shared group signal. CCA was insensitive to *σ*_indiv_ but declined with increasing noise (mean CH from 117.4 to about 81).

RoDPAA, JIVE and MOFA, by contrast, maintained strong group separation in all nine conditions. RoDPAA and JIVE were insensitive to *σ*_indiv_, with mean CH indices of 121.0, 119.0, and 115.6 for *σ*_noise_ = 1, 2, 3, respectively (Fig. 2b, Supplementary Table 1). The CH indices of RoDPAA and JIVE differed by 0.004 on average across the 900 replicates (range *−*3.9 to 2.2). This robustness is expected: stronger view-specific variation does not alter the signal subspaces or their cross-view alignment, whereas stronger noise makes the subspaces harder to estimate. MOFA, whose shared factors were identified post hoc from the variance explained in each view, had similar mean CH indices (97.8–119.5) but was more variable, with occasional low values when *σ*_indiv_*≥* 2 (Fig. 2b).

RoDPAA required about 0.14 s per run, compared with 0.73 s for MOFA and about 18 s for JIVE; CCA and the PCA-based methods required less than 0.1 s (Fig. 2c). Thus, with Gaussian data and the true ranks supplied, a setting that favours the competing methods, RoDPAA, JIVE and MOFA recovered the shared group structure similarly well, with RoDPAA the fastest.

### 2.3 RoDPAA correctly identifies the shared rank under complete and partial cross-view sharing

In practice, the shared dimension is unknown. We next asked whether it could be identified directly from the data as the degree of cross-view sharing changed. Simulation II compared RoDPAA, JIVE, and MOFA, which can estimate the shared dimension and here were required to do so; PCA and CCA were excluded because they do not estimate the dimension of a shared representation. We considered two settings in which the additional pair of latent factors was either uncorrelated across views (*ϕ* = 0) or partially aligned (*ϕ* = 0.3), corresponding to population shared dimensions of two and three, respectively, under the RoDPAA definition (Methods).

When *ϕ* = 0.3, RoDPAA selected 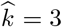 in all 100 replicates (Table 1), correctly identifying the two completely shared directions and the additional partially aligned direction with population shared intensity *σ*_3_ = 0.3. The first two recovered shared factors showed strong group separation, with a mean CH index of 119.3 (SD = 9.9). JIVE selected a joint rank of one in all 100 replicates and therefore did not recover the two exactly shared directions. MOFA identified two shared factors in 92 replicates, three in 7 and four in 1. This is consistent with its model: like JIVE, MOFA defines shared factors as common to both views, so the partially aligned direction is not part of its target (Methods). The first two MOFA shared factors separated the groups well on average (mean CH 113.0, SD = 28.0) but less consistently than those of RoDPAA (mean CH 119.3, SD = 9.9). When *ϕ* = 0, RoDPAA selected 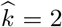 in all 100 replicates and its two shared factors separated the groups as clearly as at *ϕ* = 0.3 (mean CH 119.3, SD = 9.9). JIVE selected 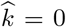 in all replicates, whereas MOFA identified two shared factors in 97 replicates and three in 3, with good but more variable group separation (mean CH 118.1, SD = 14.2). Thus, in both settings, the r.jive permutation procedure selected fewer joint components than the two exactly shared directions, which JIVE classifies as joint structure. Overall, RoDPAA recovered the population shared dimension under its definition, which includes partial sharing, in every replicate of both settings, and its group separation was the most stable.

**Table 1.**
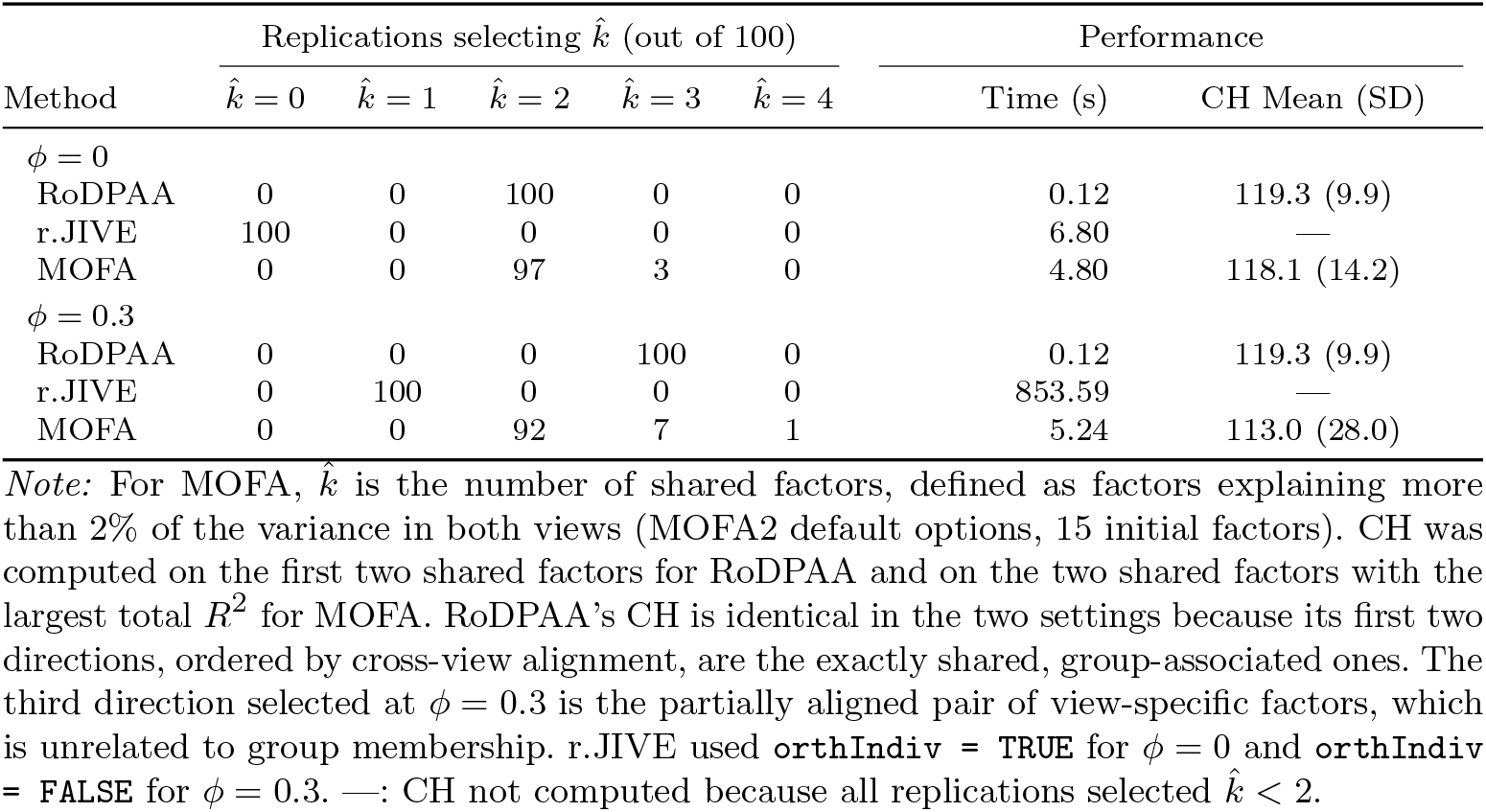
Rank selection results for three methods under *ϕ* = 0 and *ϕ* = 0.3, with *σ*_noise_ = 2, *σ*_indiv_ = 2, and 100 replications per condition. Counts show how many replications selected each rank. Time is median runtime in seconds. CH is the mean (SD) Calinski–Harabasz index conditional on 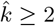.

| Method | Replications selecting k̂ (out of 100) |  |  |  |  | Performance |  |
|---|---|---|---|---|---|---|---|
|  | k̂ = 0 | k̂ = 1 | k̂ = 2 | k̂ = 3 | k̂ = 4 | Time (s) | CH Mean (SD) |
| φ = 0 |  |  |  |  |  |  |  |
| RoDPAA | 0 | 0 | 100 | 0 | 0 | 0.12 | 119.3 (9.9) |
| r.JIVE | 100 | 0 | 0 | 0 | 0 | 6.80 | — |
| MOFA | 0 | 0 | 97 | 3 | 0 | 4.80 | 118.1 (14.2) |
| φ = 0.3 |  |  |  |  |  |  |  |
| RoDPAA | 0 | 0 | 0 | 100 | 0 | 0.12 | 119.3 (9.9) |
| r.JIVE | 0 | 100 | 0 | 0 | 0 | 853.59 | — |
| MOFA | 0 | 0 | 92 | 7 | 1 | 5.24 | 113.0 (28.0) |

*Note:* For MOFA, k̂ is the number of shared factors, defined as factors explaining more than 2% of the variance in both views (MOFA2 default options, 15 initial factors). CH was computed on the first two shared factors for RoDPAA and on the two shared factors with the largest total *R*<sup>2</sup> for MOFA. RoDPAA's CH is identical in the two settings because its first two directions, ordered by cross-view alignment, are the exactly shared, group-associated ones. The third direction selected at φ = 0.3 is the partially aligned pair of view-specific factors, which is unrelated to group membership. r.JIVE used orthIndiv = TRUE for φ = 0 and orthIndiv = FALSE for φ = 0.3. —: CH not computed because all replications selected k̂ < 2.

Automatic rank estimation added little computational cost: RoDPAA, which here also estimated the signal ranks, required a median of 0.12 s per dataset in both settings, compared with 4.80 and 5.24 s for MOFA, which started from its default of 15 factors. JIVE required 6.80 s at *ϕ* = 0 and 853.59 s at *ϕ* = 0.3, depending on whether its individual subspaces could be constrained to be orthogonal (Methods).

### 2.4 Bray–Curtis-based RoDPAA yields robust shared microbiome structure across prevalence-filtering choices

Having shown that RoDPAA accurately and stably recovers shared structure under Euclidean geometry in simulation, we next examined its behavior in sparse micro-biome data, where non-Euclidean dissimilarities are routinely used [6, 7, 21]. Because prevalence-filtering thresholds are chosen by the analyst and can substantially alter the retained taxa, we asked whether the shared structure recovered by RoDPAA remained stable across prevalence-filtering choices and whether this stability depended on the underlying dissimilarity.

We analyzed data from the Inflammatory Bowel Disease Multi-omics Database (IBDMDB), a longitudinal multi-omics study of inflammatory bowel disease [3]. To investigate shared microbial structure across intestinal sites, we used paired baseline microbiome profiles from the ileum and rectum as two views of the same participants. Of the 132 enrolled participants, 68 had eligible paired baseline samples (Methods). Each anatomical site was represented by 476 genus-level microbial features. Both views were highly sparse: zeros accounted for 86.9% of ileum measurements and 87.7% of rectum measurements (Fig. 3a). Despite this, the two sites were similar at the marginal level—paired Wilcoxon tests identified only 9 of 476 genera with nominal differences (*P <* 0.05) and none after multiple-testing adjustment (Fig. 3b,c). This marginal similarity motivates an analysis of their shared multivariate structure.

**Fig. 3.**
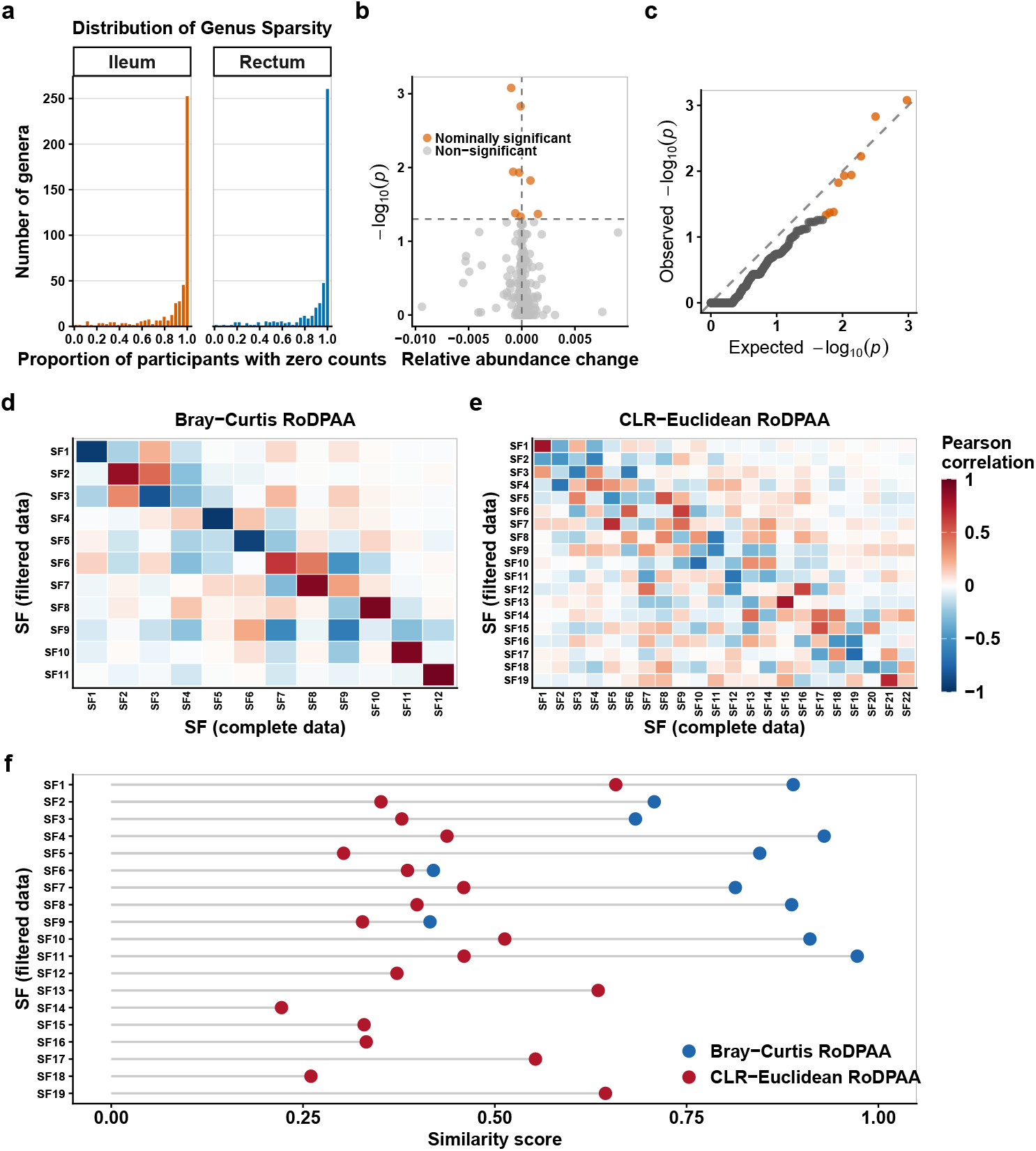
Assessment of microbiome sparsity and the sensitivity of RoDPAA shared-factor recovery to prevalence filtering. **a**, Distributions of genus-level zero proportions in the paired ileum and rectum profiles before filtering. The zero proportion is the fraction of participants in whom a feature was not detected. **b**, Mean paired relative-abundance difference between the rectum and ileum for each genus-level feature, plotted against the raw *P*-value from the paired Wilcoxon signed-rank test on the *−*log_10_ scale. Orange points indicate nominal significance at *P <* 0.05. No feature remained significant after false discovery rate correction. **c**, Quantile–quantile plot comparing the observed raw *P*-values from the 476 paired Wilcoxon tests with those expected under a uniform null distribution. **d**,**e**, Pearson correlations between shared factors (SF) estimated from prevalence-filtered and complete profiles under the Bray–Curtis (**d**) and CLR-Euclidean (**e**) configurations. Rows correspond to filtered-data shared factors, columns correspond to complete-data shared factors. **f**, Matching-concentration similarity scores for the filtered shared factors under the Bray–Curtis and CLR-Euclidean configurations. For each filtered factor, the score measures whether its correlations are concentrated on a single complete-data factor. Values near one indicate a clear one-to-one match. Together, panels d–f show greater stability of the Bray–Curtis shared factors to prevalence filtering.

We applied RoDPAA to the complete profiles and to a prevalence-filtered subset retaining genera detected in at least 20% of participants within each view (99 ileal and 94 rectal genera), using either Bray–Curtis dissimilarity or Euclidean distance after CLR transformation (hereafter Bray–Curtis-based and CLR-Euclidean RoDPAA; Methods). Under Bray–Curtis dissimilarity, the estimated shared dimension changed only slightly, from 12 to 11, whereas under CLR-Euclidean distance it decreased from 22 to 19. Because the number of components alone does not show whether the same participant-level structure was recovered, we also correlated the shared-factor scores from the filtered and complete data; under perfect stability, each filtered factor would match exactly one complete-data factor (Fig. 3d,e). Under Bray–Curtis dissimilarity, most filtered factors had one dominant complete-data counterpart (Fig. 3d and Supplementary Table 2a; maximum absolute correlation (MAC) 0.63–0.96, median 0.915, across 11 factors), so the participant-level shared structure was largely preserved even though about four-fifths of the genera were removed. CLR-Euclidean factors instead correlated diffusely with several complete-data factors (Fig. 3e and Supplementary Table 2a; MAC 0.45–0.81, median 0.614, across 19 factors). A matching-concentration score, which approaches one when a filtered factor matches a single complete-data factor (Methods), confirmed this difference (median 0.845 for Bray–Curtis versus 0.386 for CLR-Euclidean; Fig. 3f and Supplementary Table 2a). We also applied MOFA, which recovered the exactly shared dimension in most Simulation II replicates; JIVE was not applied, because its selected joint ranks in Simulation II were below the true value (Methods). MOFA, fitted with its default settings to the same CLR-transformed profiles, was less stable than CLR-Euclidean RoDPAA. It identified one shared factor in the complete data and six after filtering, and only the first filtered factor corresponded to the complete-data factor (MAC 0.69, compared with 0.01–0.18 for the other five; Supplementary Table 2a and Supplementary Fig. 1).

Together, these results highlight a key advantage of RoDPAA: the shared structure can be defined under a data geometry chosen to reflect the characteristics of the underlying modality. Here, Bray–Curtis dissimilarity made the shared structure markedly more robust to a routine preprocessing choice than the CLR-Euclidean representation used by CLR-Euclidean RoDPAA and MOFA. Based on this stability, we used Bray–Curtis dissimilarity for the microbiome view in the subsequent host–microbiome integration analyses.

### 2.5 RoDPAA with modality-specific dissimilarities reveals clinically meaningful host–microbiome structure in IBD

We next asked whether the robustness of Bray–Curtis-based RoDPAA translates into clinically meaningful discoveries in a multi-omics setting. We analyzed paired gut microbiome and metabolomic profiles from 220 participants in the Franzosa IBD study [4], including 88 participants with Crohn’s disease, 76 with ulcerative colitis, and 56 non-IBD controls (Methods). Crohn’s disease and ulcerative colitis were combined into a single IBD group. The microbiome and metabolomic views contained 11,720 genus-level microbial taxa and 8,848 metabolomic features and differed markedly in sparsity (76.3% and 27.3% zeros, respectively; Extended Data Fig. 1).

We first applied RoDPAA with a modality-specific geometry: Bray–Curtis dissimilarity for the sparse microbial profiles and Euclidean distance for the normalized metabolomic profiles (Methods). This configuration identified 30 statistically significant shared directions (Fig. 4a). In contrast, when both views were represented using Euclidean geometry, with CLR transformation applied to the microbiome data (CLR-Euclidean RoDPAA), RoDPAA identified no statistically significant shared directions (Fig. 4b), despite clear pairwise cross-omics correlations (Extended Data Fig. 2). The first two Bray–Curtis shared factors also separated participants by IBD status: unsuper-vised *k*-means clustering on these factors achieved 76.4% accuracy (adjusted Rand index (ARI) 0.267). For comparison, we extracted the two most closely aligned CLR-Euclidean directions, although neither was significant; clustering on these achieved 62.7% accuracy (ARI 0.054) (Fig. 4a,b). MOFA, fitted to the CLR-transformed microbiome and log-transformed metabolomic profiles, identified two shared factors, but these separated participants by IBD status poorly (accuracy 64.1%, ARI 0.053; Supplementary Table 2b, Supplementary Fig. 2, and Extended Data Fig. 3c). Mirroring the IBDMDB analysis, prevalence filtering left the shared dimension of Bray–Curtis-based RoDPAA unchanged (30 directions) but changed that of CLR-Euclidean RoDPAA from 0 to 67 (Extended Data Fig. 3a,b). After filtering, MOFA identified three shared factors, and its clustering accuracy fell to 56.4%, below that of filtered Bray–Curtis-based (71.4%) and CLR-Euclidean (59.1%) RoDPAA (Extended Data Fig. 3c,d; Supplementary Fig. 2; Supplementary Table 2b).

**Fig. 4.**
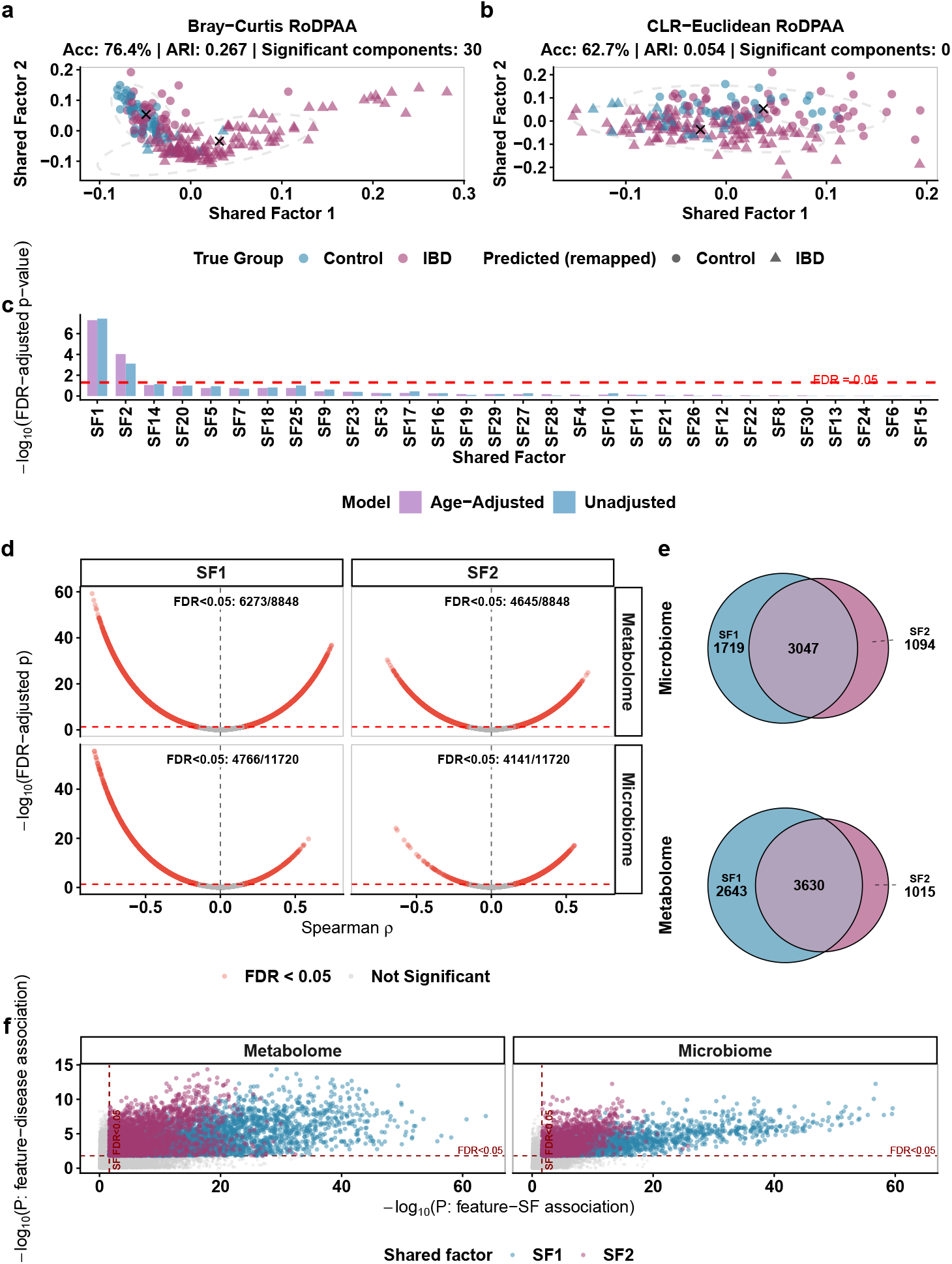
RoDPAA shared-factor representations and disease-associated feature analyses in the Franzosa IBD dataset. **a**, First two RoDPAA shared factors obtained using Bray–Curtis dissimilarity for the microbiome view and Euclidean distance for the metabolomic view. RoDPAA identified 30 statistically significant shared directions. Points are colored by true disease status and shaped by the optimally matched *k*-means cluster assignments. Ellipses represent 95% data ellipses for the estimated clusters, and crosses mark cluster centers. **b**, First two principal-angle alignment directions obtained using Euclidean distance for both views, after CLR transformation of the microbiome view (CLR-Euclidean RoDPAA). No directions were statistically significant 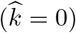 the displayed directions were extracted for exploratory visualization and clustering and are shown as in **a. c**, Associations between the 30 Bray–Curtis shared factors and IBD status from unadjusted and age-adjusted logistic regression models. Bars show *−*log_10_-transformed FDR-adjusted *P*-values, and the dashed red line indicates FDR = 0.05. **d**, Spearman associations between microbial or metabolomic features and the first two Bray–Curtis shared factors. The vertical axis shows *−*log_10_-transformed globally FDR-adjusted P-values. Red points indicate significant associations at FDR *<* 0.05, and annotations report the number of significant features. **e**, Overlap between microbial or metabolomic features significantly associated with shared factor 1 versus shared factor 2. **f**, Comparison of feature–disease and feature–shared-factor associations. The axes show *−*log_10_-transformed raw *P*-values from age-adjusted feature–disease logistic regression models and feature–shared-factor Spearman tests. Gray points represent features that were not significantly associated with disease after FDR correction; colored points represent disease-associated features, with color indicating the corresponding shared factor. Dashed red lines indicate the raw *P*-value thresholds corresponding to a global FDR of 0.05.

We then tested each of the 30 Bray–Curtis shared factors for association with IBD by logistic regression (Fig. 4c; Methods). Although RoDPAA was fitted without disease labels, the first two shared factors, denoted SF_1_ and SF_2_, were the only factors significantly associated with IBD after FDR correction, and both associations remained significant after adjustment for age. The dominant shared microbiome–metabolome variation is therefore closely related to IBD, and the clinically relevant cross-omics structure is concentrated in two dimensions.

Both factors were associated with many features in both views (Spearman correlation, FDR *<* 0.05; Fig. 4d): SF_1_ with 6,273 of 8,848 metabolomic features and 4,766 of 11,720 microbial features, and SF_2_ with 4,645 metabolomic and 4,141 microbial features. The two feature sets overlapped substantially, sharing 3,047 microbial and 3,630 metabolomic features (*χ*^2^ = 2875.4 and 249.2, both *P <* 2.2 *×*10^*−*16^; Fig. 4e), indicating overlapping cross-omics molecular programs.

Features associated with each factor were also strongly enriched for features individually associated with IBD after age adjustment (Fisher’s exact tests, all *P <* 2.2*×*10^*−*16^; Fig. 4f; Extended Data Fig. 4). This enrichment was much stronger for SF_1_ than for SF_2_ in both modalities (the odds ratios were 47.46 versus 6.92 for microbial features and 12.46 versus 2.50 for metabolomic features).

Finally, metabolites associated with SF_1_ and SF_2_ were enriched for several chemical classes annotated in the Franzosa dataset [4, 22] (hypergeometric tests, FDR *<* 0.05; Extended Data Fig. 5; Supplementary Table 3; Methods), including triterpenoids, diterpenoids and indoles for SF_1_, and carboximidic acids and tetrapyrroles for SF_2_. Cholesterols and derivatives, cholesteryl esters, and sphingolipids were enriched for both shared factors, consistent with the broad perturbations of lipid, bile-acid, and tetrapyrrole metabolism previously reported in the Franzosa IBD cohort [4].

## 3 Discussion

Multiview microbiome integration has lacked a method that operates natively in the dissimilarity space already used in single-view ecology [6, 7]. RoDPAA fills this gap: by defining shared variation through between-sample relationships within each view, it incorporates established beta-diversity dissimilarities directly, without mapping all views into a common Euclidean representation or requiring a prespecified parametric model. Our results show that this formulation matters in practice. In simulations with Gaussian data and Euclidean geometry, RoDPAA, JIVE and MOFA performed similarly when the true ranks were supplied, as expected. Defining sharing through principal-angle alignment, with the shared dimension determined by permutation inference, additionally allowed RoDPAA to recover partially shared structure. JIVE and MOFA, which define sharing as exact, are not designed to capture such directions. Because estimation requires only eigendecomposition of sample-by-sample similarity matrices and permutation, RoDPAA is also computationally efficient and needs little tuning at the modest sample sizes typical of multiview microbiome studies.

Our results also show that the well-documented sensitivity of microbiome analyses to preprocessing extends directly to multiview integration, where preprocessing can change not only the estimated shared structure but the qualitative conclusion of whether shared structure is detected at all. In the Franzosa cohort, CLR-Euclidean RoDPAA detected no shared microbiome–metabolome structure before prevalence filtering and 67 shared directions after it, whereas under Bray–Curtis dissimilarity the shared structure was stable in both cohorts. MOFA supports non-Gaussian likelihoods, but its developers recommend transforming data and using a Gaussian likelihood [23]. Applied in this way to CLR-transformed profiles, it was similarly sensitive: in the IBDMDB cohort its shared dimension changed from one to six after filtering. Because CLR-Euclidean RoDPAA was also unstable, this sensitivity is not specific to one integration method. A plausible explanation lies instead in how the two geometries handle zeros and weight rare taxa. Bray–Curtis can be applied directly to sparse abundance profiles without zero replacement or log-ratio transformation and is dominated by abundant taxa [7], so removing low-prevalence taxa changes between-sample distances little, consistent with previous evidence that prevalence filtering can largely preserve beta-diversity structure [24]. Under CLR, by contrast, every retained taxon contributes on the log scale regardless of its abundance, and zero-replaced entries become large negative values whose influence depends on the replacement value; the set of retained taxa can therefore reshape the entire geometry [12, 14]. More broadly, these findings show that RoDPAA can make multiview conclusions less dependent on a single, and often debatable, microbiome preprocessing pipeline.

The connection to beta diversity also places RoDPAA naturally within the distance-based toolkit of microbial ecology. RoDPAA starts from the same double-centered similarity matrix as PCoA [5] but asks a multiview question: which dominant axes of between-sample variation are shared across views, how strongly are they aligned, and which alignments exceed what would be expected by chance? Unlike global concordance measures such as the Mantel test or Procrustes analysis [25, 26], it estimates how many axes are shared and their shared intensities, and provides participant-level scores for each axis, enabling direct association with clinical outcomes or molecular features. In the Franzosa cohort, shared intensity itself proved informative: the two factors with the largest shared intensities were the only ones associated with IBD, and feature-level disease enrichment was substantially stronger for the first than for the second.

Several methodological questions remain open. First, inference on the shared dimension is conditional on the estimated signal subspaces. We selected RoDPAA signal ranks using a variance-explained criterion because more formal rank-selection procedures, including parallel analysis, bi-cross-validation and optimal singular-value thresholding [27–29], generally require clearer separation between signal and noise than is often present in sparse, high-dimensional microbiome data. Second, the permutation procedure assumes exchangeability of participants with respect to cross-view pairing. This assumption is appropriate for the cross-sectional settings considered here, but repeated measures, related individuals or structured batch effects would require restricted permutation schemes [30], and longitudinal designs motivate an extension that distinguishes persistent from evolving cross-view structure [3, 31]. Finally, extending RoDPAA beyond two views raises a more fundamental question about how sharing itself should be defined. Variation may be shared across all views or only subsets of them, while corresponding directions may remain only partially aligned. Existing approaches either estimate a common factor set and determine sharing post hoc, as in MOFA [19], or model subset-specific sharing explicitly, as in SLIDE [32], for which the number of possible sharing patterns grows exponentially with the number of views. A multiview extension of RoDPAA will therefore require a principled definition of partial, subset-specific sharing.

The principle underlying RoDPAA may also extend well beyond the applications considered here. Within microbiome research, the framework can accommodate other established dissimilarities, including phylogenetically informed UniFrac distances [21], and could be used to relate microbial communities across additional body sites [1, 31] or to integrate microbiome profiles with host transcriptomic, immune or metabolic measurements in other diseases [2, 33]. More generally, RoDPAA requires only a dissimilarity or similarity representation for each view, so it could, for example, identify the axes of organization shared between structural and functional brain connectivity [34, 35], or resolve the single matrix-level correlation used in representational similarity analysis [36] into multiple shared dimensions. Although these applications will require domain-specific validation, they illustrate the broader potential of RoDPAA as a computationally efficient framework for determining what different views of the same system have in common.

## 4 Methods

### 4.1 The RoDPAA framework

#### The RoDPAA model

For each view *v* = 1, 2, 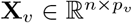 denote the observed data matrix, where the rows correspond to the same *n* subjects across the two views. We assume that **X**_*v*_ admits the decomposition

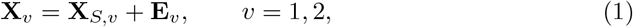

where **X**_*S,v*_ represents the underlying low-rank signal matrix and **E**_*v*_ denotes random noise. Let **K**_*S,v*_ *∈* ℝ^*n×n*^ denote the sample-by-sample similarity matrix constructed from the signal matrix **X**_*S,v*_. We assume that **K**_*S,v*_ is positive semidefinite and has rank *r*_*v*_, with spectral decomposition 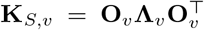, where 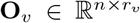 contains the eigenvectors corresponding to the *r*_*v*_ positive eigenvalues collected in 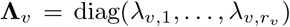. The column space *S* _*v*_ = span(**O**_*v*_) is referred to as the signal subspace of view *v*.

The overlap between the signal subspaces of the two views is characterized through the overlap between *S*_1_ = span(**O**_1_) and *S* _2_ = span(**O**_2_). Let *k* denote the number of positive singular values of 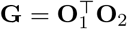. Consequently, *k* = rank(**G**) represents the dimension of the non-orthogonal overlap between the two signal spaces. Unlike existing approaches that define shared information through identical latent factors or identical subspaces, RoDPAA adopts a more flexible notion of sharing. Specifically, the shared information is represented by a pair of corresponding *k*-dimensional subspaces, one in each view, whose directions need not coincide. Instead, the degree of information sharing is quantified by their geometric alignment: directions associated with larger singular values correspond to stronger sharing, whereas smaller positive singular values indicate weaker, but still nontrivial, shared information. Therefore, the shared components are allowed to reside in distinct but geometrically related subspaces, with the singular values providing a continuous measure of the strength of information sharing.

Based on this characterization, we define the shared components as follows.

##### Definition 4.1.1

**(Shared subspace correspondence).**

Let

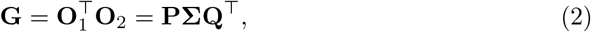

where **Σ** = diag(*σ*_1_, …, *σ*_*k*_), *k* = rank(**G**).

- **Shared subspace correspondence**. The shared information between the two views is represented by a pair of corresponding *k*-dimensional subspaces *J*_1_ = span(**O**_1_**P**) and *J*_2_ = span(**O**_2_**Q**), where each subspace belongs to its respective view. Consequently, RoDPAA allows *J*_1_ ≠ *J*_2_.
- **Paired shared directions**. For each *j* = 1, …, *k*, define **f**_1,*j*_ = **O**_1_**p**_*j*_ and **f**_2,*j*_ = **O**_2_**q**_*j*_. The pair (**f**_1,*j*_, **f**_2,*j*_) is called the *j*th *paired shared direction*. The two vectors represent the same shared latent component in the two views, but need not coincide. Their geometric alignment satisfies 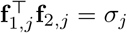, where *σ*_*j*_ *∈* (0, 1] is called the *shared intensity*. A larger value of *σ*_*j*_ indicates stronger information sharing, with *σ*_*j*_ = 1 corresponding to complete sharing and 0 *< σ*_*j*_ *<* 1 representing partial sharing.

##### The RoDPAA algorithm

Although Definition 4.1.1 provides a simple characterization of shared information through the singular value decomposition of **G** = **O**^*⊤*^_1_ **O**_2_, its direct application is infeasible in practice because the required true signals are unknown. Specifically, three practical challenges must be addressed. First, the population similarity matrices are not observed and must be estimated from the data. Second, the signal ranks *r*_1_ and *r*_2_, which determine the signal subspaces, are unknown and require data-driven estimation. Third, the signal subspaces themselves cannot be recovered by simply extracting the eigenvectors associated with the nonzero empirical eigenvalues, since finite-sample noise perturbs the empirical spectrum and obscures the boundary between signal and noise. Consequently, reliable estimation of the shared subspace correspondence requires statistically principled procedures for constructing similarity matrices, estimating the signal ranks, and identifying the signal eigenspaces in the presence of noise. RoDPAA estimates the shared structure through the following five steps.

#### Step 1. Construct similarity matrices

For each view *v* = 1, 2, compute the pairwise distance or dissimilarity matrix **D**_*v*_ = (*d*_*v,ij*_)_*n×n*_, where *d*_*v,ij*_ denotes the dissimilarity between subjects *i* and *j*, calculated from the observed data matrix **X**_*v*_ using a measure appropriate for that data type. Here, a distance is treated as a particular type of dissimilarity.

The corresponding centered sample–sample similarity matrix is constructed through Gower double-centering:

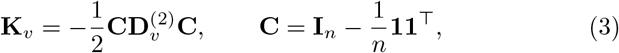

where 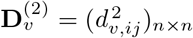 denotes the element-wise squared dissimilarity matrix.

#### Step 2. Estimate the signal subspaces

The observed similarity matrix **K**_*v*_ is a noisy version of the latent signal kernel **K**_*S,v*_. We therefore estimate the signal rank and signal subspace from the leading eigenstructure of **K**_*v*_.

First, compute the eigendecomposition

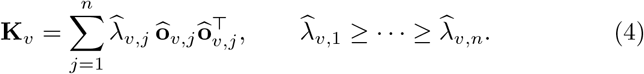

Then, estimate the signal rank 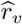, which estimates *r*_*v*_ = rank(**K**_*S,v*_), using the rank-selection procedure described in ‘RoDPAA implementation details’. Finally, retain the first 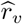 eigenvectors to form 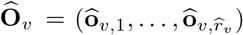, which estimates the signal subspace span(**O**_*v*_) of **K**_*S,v*_.

#### Step 3. Estimate subspace alignment

Using the estimated signal subspaces from Step 2, compute the cross-subspace matrix 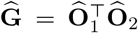, and obtain its compact singular value decomposition 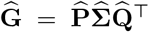, where 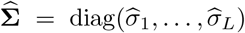 with 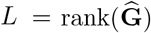, and 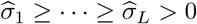.

#### Step 4. Permutation inference

The compact singular value decomposition in Step 3 produces 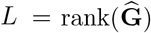 nonzero singular values. However, since the observed data matrix **X**_*v*_ is a noisy realization of the underlying signal matrix **X**_*S,v*_, some of these singular values may arise from noise rather than genuine shared information. Consequently, *L* should not be interpreted as an estimate of the shared dimension *k*. RoDPAA therefore performs statistical inference to identify the principal-angle alignments that represent significant shared information. Specifically, RoDPAA considers *H*_0*j*_ : *σ*_*j*_ = 0 versus *H*_1*j*_ : *σ*_*j*_ *>* 0 for *j* = 1, …, *L*. Under *H*_0*j*_, the *j*th principal-angle alignment does not represent systematic subject-level correspondence between the two views. To construct the permutation null distribution, RoDPAA treats the cross-view subject pairing as exchangeable and, for permutation *b*, defines 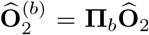, where **Π**_*b*_ is an *n × n* permutation matrix. This permutation disrupts the subject-level correspondence between the two views while preserving the orthonormality and within-view geometry of **Ô** _2_, since 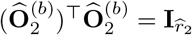.

RoDPAA then forms the permuted cross-subspace matrix 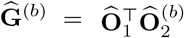 and computes its compact singular value decomposition. Let 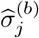 denote its *j*th singular value. The permutation *P*-value associated with the *j*th observed singular value is

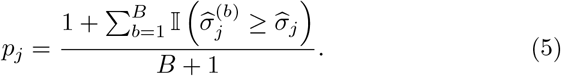

The estimated shared directions are those whose singular values are declared statistically significant, that is, whose permutation *P*-values fall below 0.05*/L* after Bonferroni correction across the *L* tested alignments.

Let 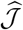. The estimated shared dimension is then 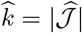.

#### Step 5. Estimate the shared directions

Following Definition 4.1.1, for each 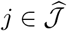, estimate the paired shared directions by 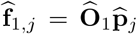 and 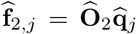, where 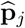and 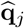are the *j*th columns of 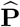 and 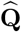, respectively. Collecting these directions gives 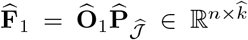 and 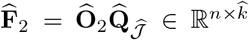. The corresponding shared intensities are 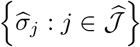. Because 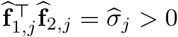, each pair is consistently oriented, and the participant-level shared factor is defined as the average of the paired directions, 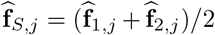. We write SF_*j*_ for the *j*th shared factor, ordered by decreasing shared intensity.

Computational details for Steps 1–5 are provided in Supplementary Algorithm 1. *Computational cost*. After the dissimilarities are computed, which requires *O*(*n*^2^*p*_*v*_) operations for view *v*, RoDPAA works only with *n × n* matrices. Step 2 requires one eigendecomposition per view, at *O*(*n*^3^) cost regardless of the number of features, and each permutation in Step 4 requires only the product 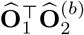, at 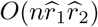cost, and the singular values of this 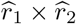 matrix, without re-estimating the signal subspaces.Apart from the signal ranks and the number of permutations, the procedure involves no iterative optimization or tuning parameters, which makes it well suited to the modest sample sizes of multiview microbiome studies.

##### RoDPAA as a generalization of JIVE and AJIVE

RoDPAA is built upon the same low-rank-plus-noise formulation adopted by many existing multiview dimension-reduction methods, including JIVE [9], one of the earliest and most influential methods for joint dimension reduction. The key distinction lies not in the statistical model, but in how shared information is defined. To illustrate this connection, we first review the JIVE formulation. Consider two data views, 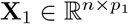 and 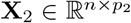, measured on the same *n* subjects. As in RoDPAA, JIVE assumes that each view admits a low-rank-plus-noise decomposition, **X**_*v*_ = **X**_*S,v*_ + **E**_*v*_, *v* = 1, 2, where **X**_*S,v*_ is the low-rank signal matrix and **E**_*v*_ is the residual noise matrix.

JIVE further decomposes the signal matrix as

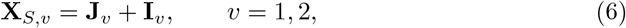

where **J**_*v*_ and **I**_*v*_ denote the joint and view-specific individual signal components, respectively. To ensure identifiability of the decomposition, JIVE further assumes that

1. the joint components share the same subject-level subspace, col(**J**_1_) = col(**J**_2_) = *S*_*J*_, where *S*_*J*_ *⊆* ℝ^*n*^;
2. for each view, col(**I**_*v*_) ⊥ *S*_*J*_;
3. the individual components satisfy col(**I**_1_) *∩* col(**I**_2_) = *{***0***}*.

Under these assumptions, JIVE estimates the joint and individual components by minimizing the reconstruction error

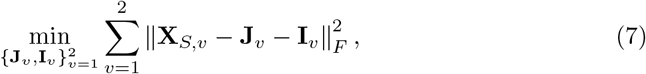

subject to the rank and identifiability constraints. The resulting alternating optimization repeatedly updates the joint and individual components until convergence. Although statistically well motivated, the repeated low-rank approximations required by this iterative procedure may become computationally expensive for high-dimensional data. To improve computational efficiency, Angle-based JIVE (AJIVE) was proposed as a subspace-based reformulation of JIVE [10]. Rather than iteratively estimating the joint and individual matrices, AJIVE first estimates the signal subspace for each view and then identifies the joint subspace through principal-angle analysis. Importantly, AJIVE retains the JIVE definition of shared information: the joint component is characterized by an identical subject-level subspace across the two views.

Under the JIVE model, the population joint subspace corresponds to zero principal angles, or equivalently, principal-angle singular values equal to one. Because the signal subspaces are estimated from noisy data, the observed principal angles are generally nonzero even when the underlying population subspaces are identical. To account for this estimation error, AJIVE uses Wedin’s perturbation theorem to derive an upper bound on the rotation of the estimated signal subspaces attributable to noise. Principal angles below this perturbation bound are regarded as consistent with an underlying common subspace and are retained as joint directions. The remaining variation is obtained by orthogonally projecting each signal subspace onto the complement of the estimated joint subspace to obtain the individual components.

RoDPAA extends the JIVE/AJIVE framework along two complementary dimensions: the definition of shared structure across views and the geometry used to represent variation within each view. First, RoDPAA generalizes the definition of shared structure. Classical JIVE defines shared information through an identical subject-level latent sub-space across views (equation (6), Assumption 1). Consequently, latent directions that are strongly related across views but do not belong to exactly the same subspace are not included in the joint component. RoDPAA replaces this exact common-subspace requirement with a statistical definition of sharing based on principal-angle alignment. Corresponding directions are considered shared when their observed alignment is significantly stronger than expected under random subject correspondence, as determined by the permutation procedure in Step 4 of the RoDPAA algorithm. Exact common directions are therefore retained as a special case, while partially aligned directions can also be identified as shared when supported by the data. Second, RoDPAA generalizes the geometry used to represent each view. JIVE and AJIVE construct their signal subspaces directly from the observed data matrices, which is equivalent to using the centered linear kernel and therefore inherits the Euclidean geometry of the original feature space (Supplementary Note 1). RoDPAA instead constructs each signal subspace from a sample–sample similarity matrix derived from a user-specified dissimilarity. This allows each view to retain a data-appropriate notion of between-sample variation, including non-Euclidean dissimilarities such as Bray–Curtis, while leaving the subsequent principal-angle analysis unchanged. When squared Euclidean distance is used, Gower double centering recovers the centered linear kernel. RoDPAA therefore yields the same sample-level signal subspaces as JIVE and AJIVE under Euclidean geometry (Supplementary Note 1). Combined with exact cross-view alignment, the classical JIVE shared-subspace formulation is recovered as a special case of RoDPAA. Even in this special case, the estimators differ. JIVE obtains its joint component by iteratively fitting a low-rank approximation to the concatenated data, a variance-driven step in which strong view-specific variation can compete with the shared signal. RoDPAA estimates each signal subspace once by eigendecomposition and ranks the paired directions by their cross-view alignment. With correctly specified ranks, the two estimators nevertheless gave virtually identical results in Simulation I (Results).

### 4.2 Simulation studies

#### Data generation

For each subject *i* = 1, …, *n*, a binary group label *y*_*i*_ *∈ {−*1, +1*}* was generated with equal probability. Conditional on *y*_*i*_, we generated 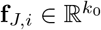 according to

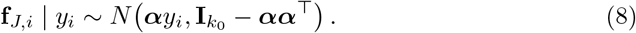

Here, 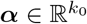 determines the magnitude of group separation through the conditional means E(**f**_*J,i*_ | *y*_*i*_ = *±*1) = *±****α***, and satisfies the constraint ∥α∥ *<* 1, ensuring that 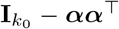is positive definite and therefore defines a valid covariance matrix.

Equivalently, **f**_*J,i*_ satisfies the two-component Gaussian mixture 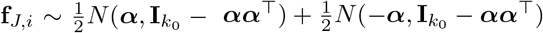. Its marginal mean and covariance are E(**f**_*J,i*_) = **0** and 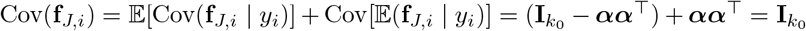. Thus, **f**_*J,i*_, *i* = 1, …, *n*, are marginally independently and identically distributed.

For the additional components, we generate one latent factor for each view with target population correlation *ϕ∈* (*−* 1, 1). To reduce finite-sample variation, we further construct the factors so that their empirical correlation is exactly *ϕ* in each simulated data set. Specifically, generate two independent Gaussian vectors 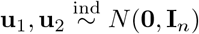.

Center and standardize **u**_1_ to obtain the additional factor for view 1, **F**_*I*,1_ *∈* ℝ^*n*^. Next, remove from **u**_2_ its component along 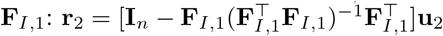. After centering and standardizing **r**_2_, denote the resulting vector by 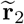, and define the additional factor for view 2 as 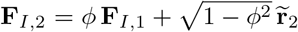. Because **F**_*I*,1_ and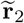 are empirically orthogonal and have unit sample variance, this construction ensures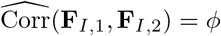.

Stacking the subject-level quantities **f**_*J,i*_ across subjects to form 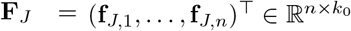, we generated the paired data matrices according to

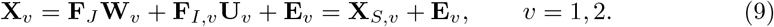

where 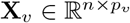 is the observed data matrix, *p*_*v*_ is the number of features in view *v*, **X**_*S,v*_ = **F**_*J*_ **W**_*v*_ +**F**_*I,v*_**U**_*v*_ represents the low-rank signal matrix described in equation (1), 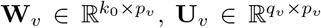, and 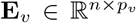is the Gaussian noise matrix. The entries of **W**_*v*_ were generated independently from *N* (0, 1), whereas the entries of **U**_*v*_ were generated independently from 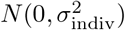. The entries of **E**_*v*_ were generated independently from 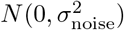. Finally, we set *n* = 300, *p*_1_ = 100 and *p*_2_ = 150, with *q*_1_ = *q*_2_ = 1 additional factor per view.

Under this design, **F**_*J*_ contains *k*_0_ = 2 latent factors common to both views, generating two completely shared directions with population shared intensities *σ*_1_ = *σ*_2_ = 1. The shared intensity associated with the directions represented by **F**_*I*,1_ and **F**_*I*,2_ is determined by the correlation parameter *ϕ*. Specifically, when *ϕ* = 0, col(**F**_*I*,1_) ⊥ col(**F**_*I*,2_), so the third pair of latent directions is orthogonal across views and does not contribute to the shared structure. The cross-view alignment spectrum is (*σ*_1_, *σ*_2_, *σ*_3_) = (1, 1, 0), with only the first two directions classified as shared. Hence, RoDPAA’s population shared dimension under Definition 4.1.1 is *k* = 2.

When *ϕ* ≠ 0, col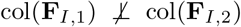, so these directions are partially shared. The corresponding shared intensity is the cosine of their principal angle: 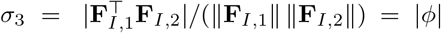|. Therefore, the population shared intensities are (*σ*_1_, *σ*_2_, *σ*_3_) = (1, 1, |*ϕ* |). For 0 *<* |*ϕ* |*<* 1, all three directions are shared under the RoDPAA definition, giving a population shared dimension of *k* = 3. In contrast, the third direction is excluded from the joint subspace under classical JIVE because it is not exactly shared (*σ*_3_ *<* 1). The same holds for MOFA, whose shared factors are common to both views by construction, so that under the JIVE and MOFA models the population shared dimension is two whether *ϕ* = 0 or *ϕ* = 0.3 (Simulation II).

##### Benchmark methods and common implementation

RoDPAA was compared with JIVE [9, 37], MOFA [19], canonical correlation analysis (CCA) [8], and principal component analysis (PCA) [38]. All methods were fitted without access to the simulated group labels. JIVE and MOFA provide direct multiview latent-factor comparisons, CCA represents a correlation-based integration approach, and PCA provides non-integrative baselines based on either individual views or their concatenation.

JIVE was implemented using the `r.jive` package, MOFA using `MOFA2` with a Gaussian likelihood and `fast` convergence mode, CCA using `CCA::cc,` and PCA using `prcomp.` PCA was applied separately to each view and to the horizontally concatenated data matrix [**X**_1_ |**X**_2_]. Because JIVE, CCA, and PCA operate on Euclidean feature representations, RoDPAA was implemented using Euclidean distance in the simulations.

This design isolates differences in recovery of shared structure rather than differences arising from the choice of dissimilarity.

##### Construction of low-dimensional representations in the simulation

For comparisons based on participant-level structure, each method was represented by its leading two components. For methods producing a single set of participant scores, including MOFA and PCA, the first two component scores were used directly; for MOFA, these were the first two shared factors. For methods producing paired view-specific scores, including RoDPAA, JIVE, and CCA, corresponding scores from the two views were combined by averaging to obtain a single participant-level representation. Specifically, if **f**_1,*j*_ and **f**_2,*j*_ denote the paired scores for component *j*, the combined score was 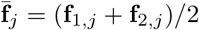 for *j* = 1, 2. The two-dimensional representation was therefore 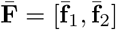. For paired components whose signs were not intrinsically aligned by the estimation procedure, the orientation of the second-view score was chosen so that its correlation with the corresponding first-view score was positive before averaging.

##### Evaluation of group separation in simulation

Because the simulated group labels were known, recovery of the group-associated latent structure was evaluated directly from the two-dimensional participant representations without performing clustering. For each method, group separation was quantified using the Calinski–Harabasz (CH) index with respect to the simulated group labels.

Specifically, recall *y*_*i*_ *∈ {−*1, 1*}* denotes the group label for subject *i*, and define *ℐ*_1_ = *{i* : *y*_*i*_ = *−*1*}* and ℐ*I*_2_ = *{i* : *y*_*i*_ = 1*}*, with *n*_*g*_ = | *ℐ*_*g*_|, *g* = 1, 2. For a recovered two-dimensional representation **f**_*i*_ *∈* ℝ^2^ of subject *i*, define the group centroid and overall centroid as 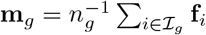and 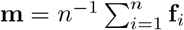. The between-group and within-group sums of squares are

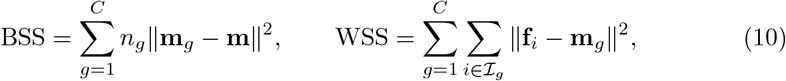

where *C* = 2 is the number of groups and *n* = 300 is the total sample size. The Calinski–Harabasz (CH) index [39] is then defined as

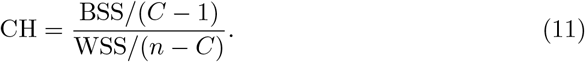

Larger CH values indicate better recovery of the group-discriminating latent structure.

##### Simulation I: Fixed-rank recovery across noise and view-specific variation

Simulation I compared recovery of the two-dimensional group-associated shared structure when the target dimension was prespecified. For RoDPAA, the signal ranks were fixed at *r*_1_ = *r*_2_ = 3, and the first two paired shared directions were retained. JIVE was fitted with `method=“given”`, joint rank two, and individual rank one (signal rank three) for each view. MOFA was fitted with four latent factors, the number of dis-tinct latent factors in the simulation model (two shared factors and one view-specific factor per view); a factor was treated as shared when it explained more than 2% of the variance in both views 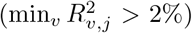. and the first two shared factors were evaluated; CCA retained the first two pairs of canonical variates, and PCA retained the first two principal components from each corresponding analysis. Measurement noise and view-specific variation were varied independently over *σ*_noise_ *∈ {*1, 2, 3*}* and *σ*_indiv_ *∈ {*1, 2, 3*}*, yielding nine conditions, with *ϕ* = 0.3. One hundred independent datasets were generated per condition, for a total of 900 replicates. For each replicate, we recorded the CH index and computation time of each method.

Simulation II evaluated whether the dimension of the shared structure could be identified directly from the data. Measurement noise and view-specific variation were fixed at *σ*_noise_ = *σ*_indiv_ = 2, and two levels of cross-view alignment were considered.

When *ϕ* = 0, the population shared dimension was two; when *ϕ* = 0.3, the partially aligned third direction increased the population shared dimension to three under the RoDPAA definition. One hundred independent datasets were generated for each setting. RoDPAA estimated the shared dimension using its permutation-based shared alignment test with Bonferroni correction, after estimating the signal ranks with an elbow criterion, that is, at the largest relative drop between consecutive positive eigenvalues of **K**_*v*_. JIVE estimated the joint rank using the permutation procedure implemented in `r.jive (method=“perm”)`. Mutually orthogonal individual subspaces were imposed (`orthIndiv=TRUE`) when *ϕ* = 0 and not imposed (`orthIndiv=FALSE`) when *ϕ* = 0.3, matching the simulated individual structure; the orthogonality constraintis stronger than JIVE’s identifiability assumptions require and substantially reduces computation time. MOFA was fitted with the MOFA2 default options (15 initial factors for this sample size, with automatic relevance determination priors on the weights), so that, like RoDPAA and JIVE, it estimated the number of factors from the data. Adapting the 2% variance criterion of the original MOFA study [19], a factor was classified as shared when it explained more than 2% of the variance in both views 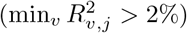, and the two shared factors explaining the most total variance were used for the CH index. PCA and CCA were not included because they do not provide an explicit estimate of the dimension of a shared latent representation. For each replicate, we recorded the estimated shared dimension and computation time. When at least two shared factors were recovered, the first two were additionally evaluated using the CH index described above.

### 4.3 Data sets and preprocessing

RoDPAA was applied to two IBD cohorts: paired ileum and rectum microbiome profiles from the IBDMDB, and paired gut microbiome and metabolome profiles from the Franzosa IBD study.

#### IBDMDB cohort and participant selection

Microbiome data were obtained from the Inflammatory Bowel Disease Multi-omics Database (IBDMDB), established through the Integrative Human Microbiome Project [3]. The full IBDMDB cohort comprised 132 participants with Crohn’s disease, ulcerative colitis, or no IBD and included longitudinal multi-omics measurements, including 16S rRNA gene sequencing of intestinal biopsy samples. Because paired site-specific ileum and rectum profiles were available for only a subset of participants, the present analysis was restricted to a baseline multi-omics cohort of 68 participants with both profiles, including 47 participants with IBD and 21 non-IBD controls.

#### IBDMDB microbiome processing and feature representation

The ileum and rectum microbiome profiles were derived from 16S rRNA gene amplicon sequencing of intestinal biopsy samples in the IBDMDB study [3]. In the original IBDMDB processing pipeline, the V4 region of the 16S rRNA gene was sequenced using the Illumina MiSeq platform, reads were demultiplexed and merged using USEARCH, and sequences were clustered into operational taxonomic units (OTUs) at 97% sequence similarity using UPARSE algorithm. Taxonomic assignments were obtained by mapping OTUs to a V4-region reference derived from the SILVA database. For the present analysis, the ileum and rectum profiles were represented by 476 genus-level microbial features and treated as two paired views of the same 68 participants.

#### Franzosa IBD study population and multi-omics data

Data were obtained from the cross-sectional Franzosa IBD study, which profiled stool samples from participants with Crohn’s disease, ulcerative colitis, and non-IBD controls using shotgun metagenomic sequencing and untargeted liquid chromatography–mass spectrometry metabolomics [4]. The study included discovery and validation cohorts of 155 and 65 participants, respectively. Microbial genus-level relative abundances were obtained from a curated multi-omics resource [40], which reprocessed the raw shotgun metagenomic reads from this study using Kraken2/Bracken against the Genome Taxonomy Database and aggregated species-level taxa to the genus level, yielding 11,720 genus-level taxa with abundances proportionally normalized to sum to one per sample. Metabolomic feature intensities, as provided by the original study, were sumnormalized to parts per million within each of four LC-MS methods (C8-pos, C18-neg, HILIC-pos, HILIC-neg) [4], yielding 8,848 features. The two cohorts were combined without additional batch-effect correction, and Crohn’s disease and ulcerative colitis were combined into a single IBD group for comparison with non-IBD controls. All available features were retained in the primary analysis. Feature filtering was considered separately as a sensitivity analysis.

#### Feature sparsity and prevalence filtering

For a data view *v*, let *X*_*igv*_ denote the observed abundance or intensity for participant *i* and feature *g*. Feature prevalence was summarized through the zero proportion 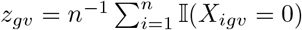, where *n* is the number of participants in the corresponding analysis. Larger values of *z*_*gv*_ indicate lower feature prevalence. Overall sparsity within each view was additionally summarized as the proportion of all participant–feature entries equal to zero. When prevalence filtering was used, it was performed separately within each view. A feature was retained when *z*_*gv*_*≤ τ*_*v*_, where *τ*_*v*_ denotes the prespecified maximum zero proportion for that analysis. Equivalently, retained features were required to be observed in at least 1 *−τ*_*v*_ of participants. The threshold used in the applications is given in ‘RoDPAA implementation details’.

### 4.4 RoDPAA implementation in the IBD applications

#### Construction of microbiome kernels

For microbiome data, the primary representation used Bray–Curtis dissimilarity applied directly to the observed abundance profiles. For participants *i* and *i*^*′*^ in view *v*,

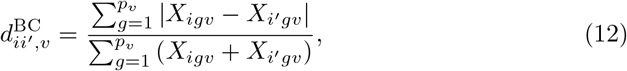

where *p*_*v*_ denotes the number of microbial features in view *v*. For comparison, we also considered a CLR-based Euclidean representation (CLR-Euclidean RoDPAA). After adding an application-specific pseudocount *δ*_*v*_ *>* 0, microbial abundances were transformed as

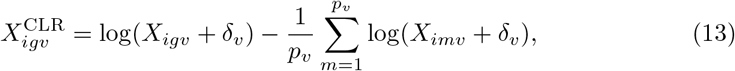

and pairwise Euclidean distances were calculated between the transformed participant profiles. For either representation, the resulting dissimilarity matrix **D**_*v*_ was converted to a centered sample–sample similarity matrix using Gower double centering (equation (3)).

#### Construction of the metabolomic kernel in the Franzosa cohort

For the metabolomic view in the Franzosa cohort, the PPM-normalized metabolome intensities described above were log-transformed before constructing Euclidean distances. Let *Z*_*il*_ denote the observed intensity of metabolomic feature *l* for participant *i*. To accommodate zero values, we added a pseudocount equal to one half of the smallest positive intensity, 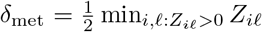, and defined 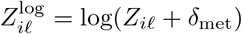. Pair-wise Euclidean distances were then calculated between the log-transformed participant profiles. The resulting distance matrix was converted to a centered sample–sample similarity matrix using the same Gower double-centering transformation described above (equation (3)).

#### RoDPAA implementation details

For each view *v*, RoDPAA was applied to the resulting centered similarity matrix **K**_*v*_. Because Gower double centering of a non-Euclidean dissimilarity such as Bray–Curtis can yield an indefinite similarity matrix, signal-subspace estimation was restricted to eigenvectors associated with positive eigenvalues. The signal rank was selected as the smallest number of positive eigenvalues explaining at least 80% of the total positive eigenvalue sum. Shared directions were then identified using the RoDPAA permutation procedure with 1,000 permutations and the significance criterion described in Step 4 of the RoDPAA algorithm. The same settings were used in the IBDMDB and Franzosa analyses.

To assess the robustness of RoDPAA to the removal of low-prevalence features, each dissimilarity configuration was fitted to both the unfiltered and the prevalence-filtered data. In the IBDMDB analysis, features detected in at least 20% of participants within each view were retained (*z*_*gv*_ *≤* 0.8). In the Franzosa analysis features detected in more than 80% of participants within each view were retained (*z*_*gv*_ *<* 0.2).

#### Comparison with MOFA in the IBD applications

MOFA was fitted with the `MOFA2` default options (15 initial factors, Gaussian likelihood, automatic relevance determination priors on the weights, at most 1,000 iterations). Although MOFA also supports Poisson and Bernoulli likelihoods [19], the `MOFA2` documentation recommends transforming such data and using the Gaussian likelihood [23]; following this recommendation, MOFA was applied to the representations used for the Euclidean RoDPAA configurations: CLR-transformed genus profiles (equation (13)) for the rectum and ileum in the IBDMDB cohort, and CLR-transformed microbiome and log-transformed metabolomic profiles in the Franzosa cohort, each centered by feature. It was therefore not applied to the Bray–Curtis representation. As in Simulation II, a factor was classified as shared when it explained more than 2% of the variance in both views, and shared factors were ordered by total variance explained. In both cohorts, MOFA was fitted to the complete and the prevalence-filtered profiles. In the IBDMDB cohort, the stability of its shared factors was assessed with the maximum absolute correlation used for RoDPAA, and a second fit to the complete profiles with a different random initialization was used to separate the effect of filtering from that of initialization. In the Franzosa cohort, *k*-means clustering on the first two shared factors was performed as for RoDPAA. JIVE was not applied to the IBD data, because in Simulation II the joint ranks selected by the `r.jive` permutation procedure were below the number of exactly shared directions, so its estimated joint structure could not be used to define shared factors.

### 4.5 Downstream statistical analyses

#### Marginal comparison between ileum and rectum in the IBDMDB cohort

To compare genus-level abundances between the two anatomical sites, raw counts were first converted to relative abundances within each sample, 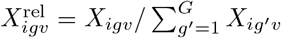, with *G* = 476. For each participant and genus, we then calculated the paired difference 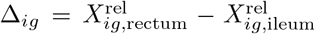 and summarized the direction and magnitude of the marginal difference by 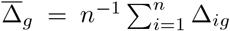 Given the extreme sparsity and non-Gaussian distribution of the microbial abundances, paired Wilcoxon signed-rank tests were used to test for genus-specific differences between the ileum and rectum. Features with raw *P <* 0.05 were highlighted as nominally significant for visualization, and the Benjamini–Hochberg procedure was applied across the 476 tests to control the false discovery rate at 0.05. The distribution of the raw Wilcoxon *P*-values was additionally examined using a quantile–quantile plot. The ordered observed *P*-values were compared with their expected order statistics under a Uniform(0, 1) null distribution, with both axes displayed on the *−* log_10_ scale.

#### Assessment of shared-factor stability in the IBDMDB application

For each dissimilarity configuration, shared factors estimated from the filtered and complete data were compared using pairwise Pearson correlations, 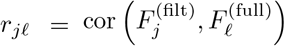. Because factor signs are arbitrary, absolute correlations were used for numerical summaries. For each filtered factor *j*, the strength of its closest match in the complete-data analysis was defined as *M*_*j*_ = max_*l*|_ *r*_*j*|*l*_.

We also defined a matching-concentration score,

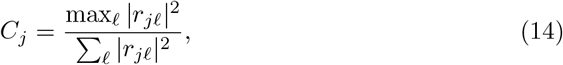

which measures whether a filtered factor corresponds primarily to a single complete-data factor or is distributed across multiple factors. Larger values indicate a clearer one-to-one match. These metrics were compared between the Bray–Curtis and CLR-based Euclidean analyses to assess stability to prevalence filtering.

#### Clustering of Franzosa participants and concordance with IBD status

For each dissimilarity configuration, *k*-means clustering was performed using the first two participant-level RoDPAA component scores, obtained by averaging paired directions as in Step 5 of the RoDPAA algorithm. When statistically significant shared directions were identified, these scores corresponded to the first two estimated shared factors. When the estimated shared dimension was zero, as in the CLR-based Euclidean analysis, the first two candidate principal-angle alignment directions were used only for exploratory visualization and clustering and were not interpreted as shared factors. The number of clusters was fixed at *K* = 2 to permit comparison with the two observed clinical groups, IBD and non-IBD control. Clustering was performed using `stats::kmeans` with 50 random initializations, retaining the solution with the smallest within-cluster sum of squares. RoDPAA estimation and cluster assignment were entirely unsupervised; disease status was used only afterward to evaluate concordance between the estimated clusters and the observed clinical groups. Agreement was quantified using permutation-invariant accuracy [41] and the adjusted Rand index (ARI) [42]. Because cluster labels are arbitrary, accuracy was calculated after selecting the mapping of estimated cluster labels to disease groups that maximized agreement,

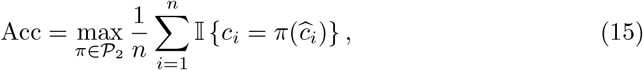

where *c*_*i*_ and 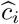 denote the observed disease group and estimated cluster label, respectively. The ARI additionally measures agreement between the two partitions while correcting for agreement expected by chance and was calculated using `mclust::adjustedRandIndex` [43].

#### Downstream association analyses in the Franzosa cohort

To evaluate the clinical relevance of the recovered shared structure, each statistically significant shared factor was tested for association with IBD status using logistic regression, both without covariate adjustment and with adjustment for age. Shared factors significant in the age-adjusted analysis were carried forward for feature-level interpretation. For each selected shared factor, Spearman correlations were calculated with all microbial and metabolomic features. Features in both domains were transformed as log(*x* + *c*), where *c* was defined as one-half of the minimum nonzero value in the corresponding data domain. For these feature-level analyses, microbial abundances were log-transformed rather than CLR-transformed because each feature was analyzed separately; CLR transformation was reserved for the multivariate Euclidean RoDPAA analysis, where between-subject dissimilarities were defined using the full microbial composition. Each microbial and metabolomic feature was also tested directly for association with IBD using age-adjusted logistic regression. The same log-transformed features were standardized to mean zero and unit variance before model fitting. For each family of tests, *P*-values were adjusted using the Benjamini–Hochberg procedure, with FDR *<* 0.05 considered significant. Adjustment was performed separately for the unadjusted and age-adjusted shared-factor models, jointly across all feature–factor tests, and jointly across all feature–IBD tests.

All overlap analyses used 2 *×* 2 contingency tables, separately for the microbial and metabolomic views. Overlap between the SF_1_- and SF_2_-associated feature sets was tested using Pearson’s *χ*^2^ test. Enrichment of features associated with each shared factor among features directly associated with IBD was tested using Fisher’s exact test and summarized by the odds ratio.

#### Metabolite chemical-class over-representation analysis

Metabolite chemical-class annotations were obtained from the metadata provided with the Franzosa study [4]. Metabolites without a Putative Chemical Class annotation were excluded from enrichment analyses. Among the annotated metabolites, chemical classes represented by at least 10 features were tested for over-representation among metabolites significantly associated with SF_1_ or SF_2_ using one-sided hypergeometric tests. Of the 8,848 metabolomic features, 3,820 had a class annotation and formed the enrichment background; of these, 2,854 and 2,157 were associated with SF_1_ and SF_2_, respectively, and 97 classes met the minimum-size criterion. Enrichment analyses were performed separately for the two shared factors, and *P*-values were adjusted across tested chemical classes using the Benjamini–Hochberg procedure; classes with *q <* 0.05 were considered significantly enriched. To characterize the overall enrichment pattern beyond the individually significant classes, observed enrichment *P*-values were also compared with their expected distribution under the uniform null using quantile–quantile plots, separately for SF_1_ and SF_2_.

## Data Availability

Processed microbial relative-abundance profiles and metabolomic feature intensities for the Franzosa IBD cohort [4] were obtained from the curated gut microbiome–metabolome data resource described by Muller et al. [40], publicly available at https://github.com/borenstein-lab/microbiome-metabolome-curated-data (dataset FRANZOSA IBD 2019, directly accessible at https://github.com/borenstein-lab/microbiome-metabolome-curated-data/tree/main/data/processed-data/FRANZOSA_IBD_2019).

Paired ileum and rectum 16S rRNA microbiome profiles for the IBDMDB cohort were obtained through the Integrative Human Microbiome Project (iHMP) [3], publicly available from the IBD Multi-omics Database at https://ibdmdb.org/results. Merged 16S taxonomic profiles are available at https://ibdmdb.org/downloads/html/products_16s_2018-01-08.html; samples were assigned to ileum or rectum biopsy sites using the accompanying HMP2 metadata file.

## Code Availability

RoDPAA software (for R version 3.6.0 or later) is available at https://github.com/yanyanxu0428/RoDPAA.

## Figures

**Extended Data Fig. 1.**
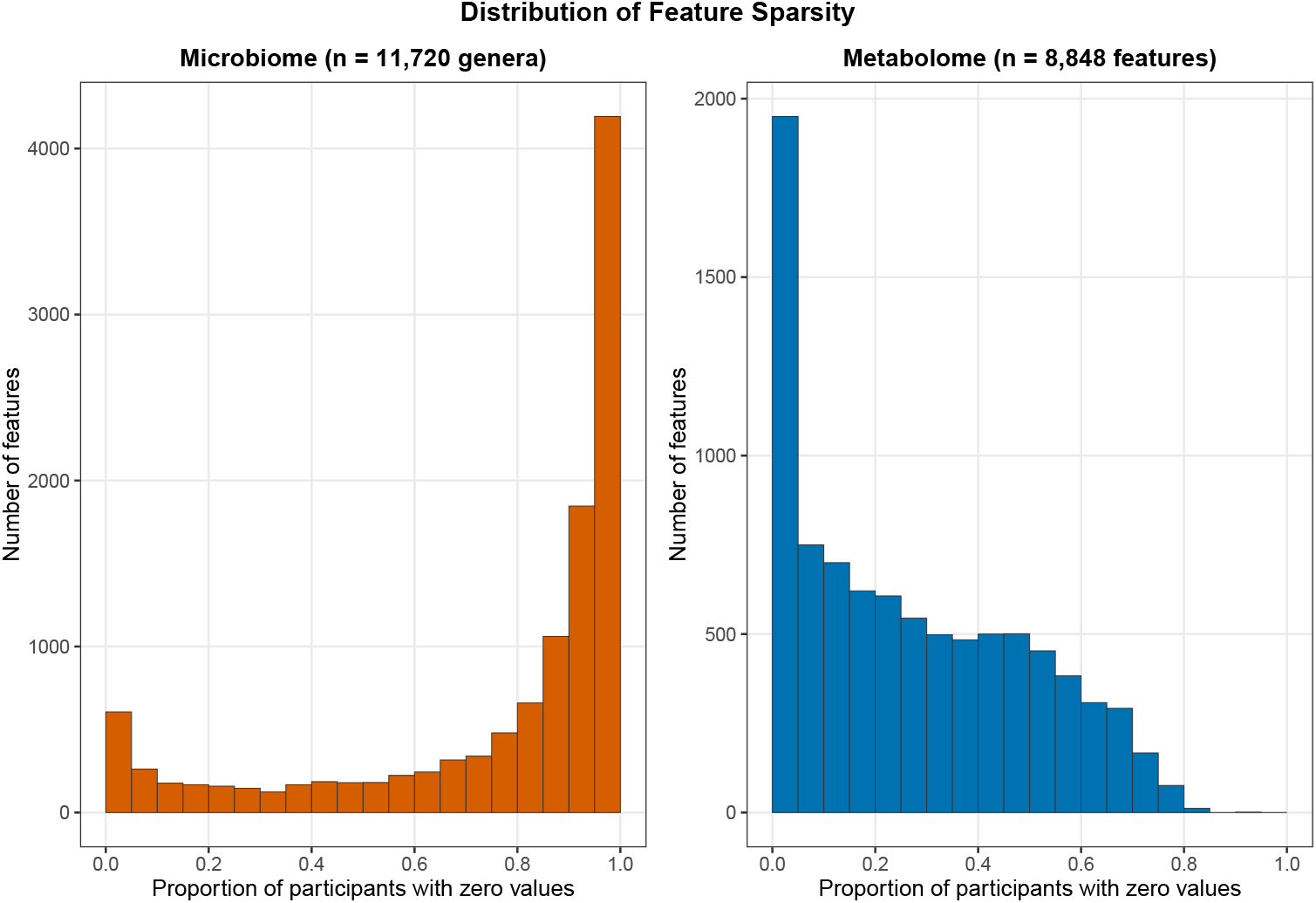
Zero proportions in the Franzosa IBD microbiome and metabolome data. Histograms of the proportion of the 220 participants with a zero value for each feature, shown for the 11,720 microbial genera (left) and the 8,848 metabolomic features (right), before any feature filtering. Overall, 76.3% of microbial and 27.3% of metabolomic values were zero. Most genera were absent from most participants (median per-genus zero proportion, 0.91; 7,760 genera, 66.2%, were zero in more than 80% of participants), whereas most metabolomic features were detected in most participants (median per-feature zero proportion, 0.24; 13 features were zero in more than 80% of participants).

**Extended Data Fig. 2.**
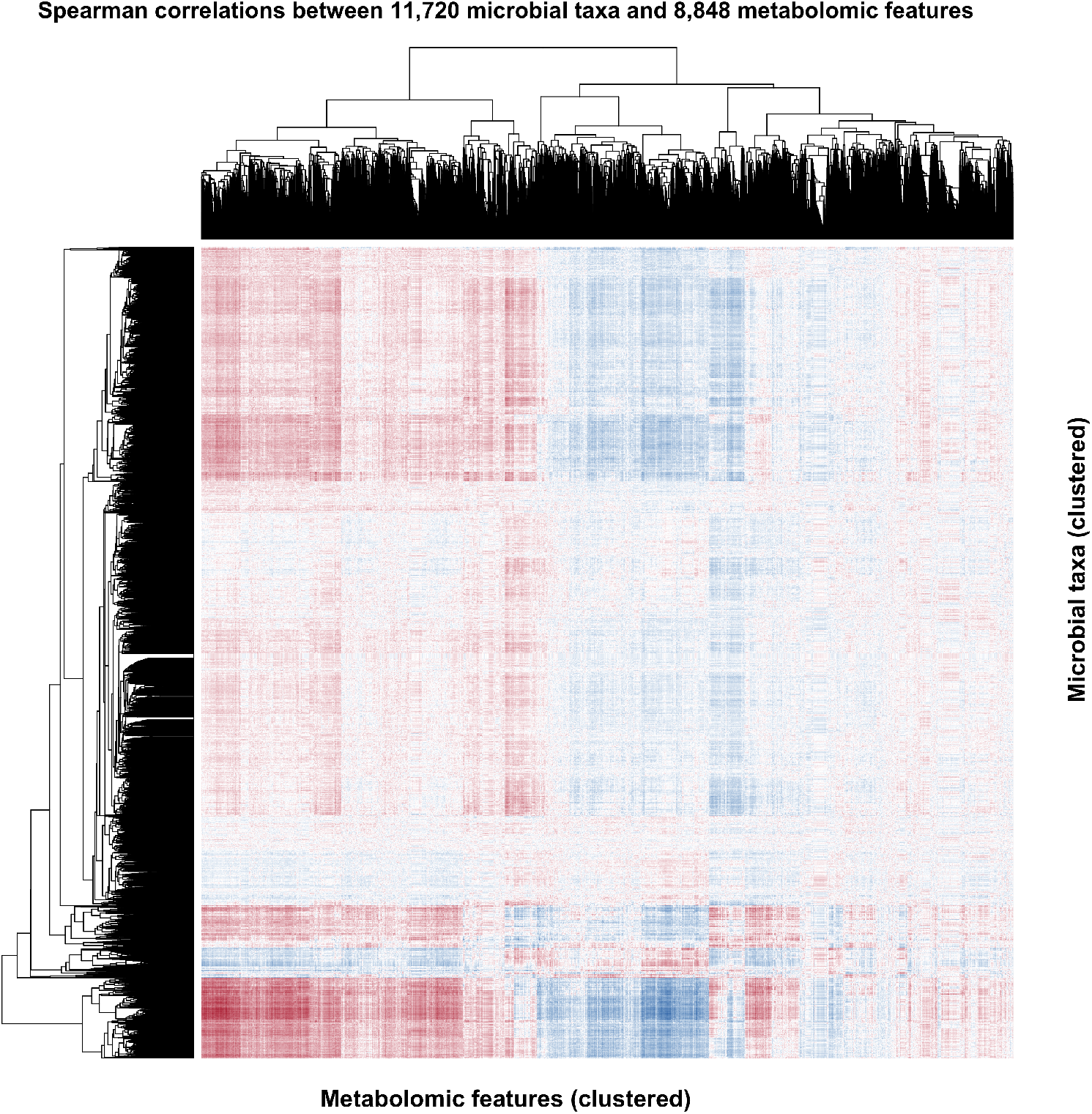
Full pairwise Spearman correlation between microbial taxa and metabolomic features in the Franzosa IBD cohort. Heatmap of Spearman correlation coefficients between all 11,720 microbial taxa (rows) and all 8,848 metabolomic features (columns) across 220 paired samples. Both axes are ordered by average-linkage hierarchical clustering (dendrograms shown), computed on Euclidean distances between each feature’s correlation profile. Color indicates the Spearman correlation coefficient (dark red, strong positive; dark blue, strong negative; white, no correlation). Large contiguous blocks of concordant color indicate that groups of taxa and groups of metabolites share coordinated cross-omics correlation structure, rather than correlations being distributed at random across feature pairs.

**Extended Data Fig. 3.**
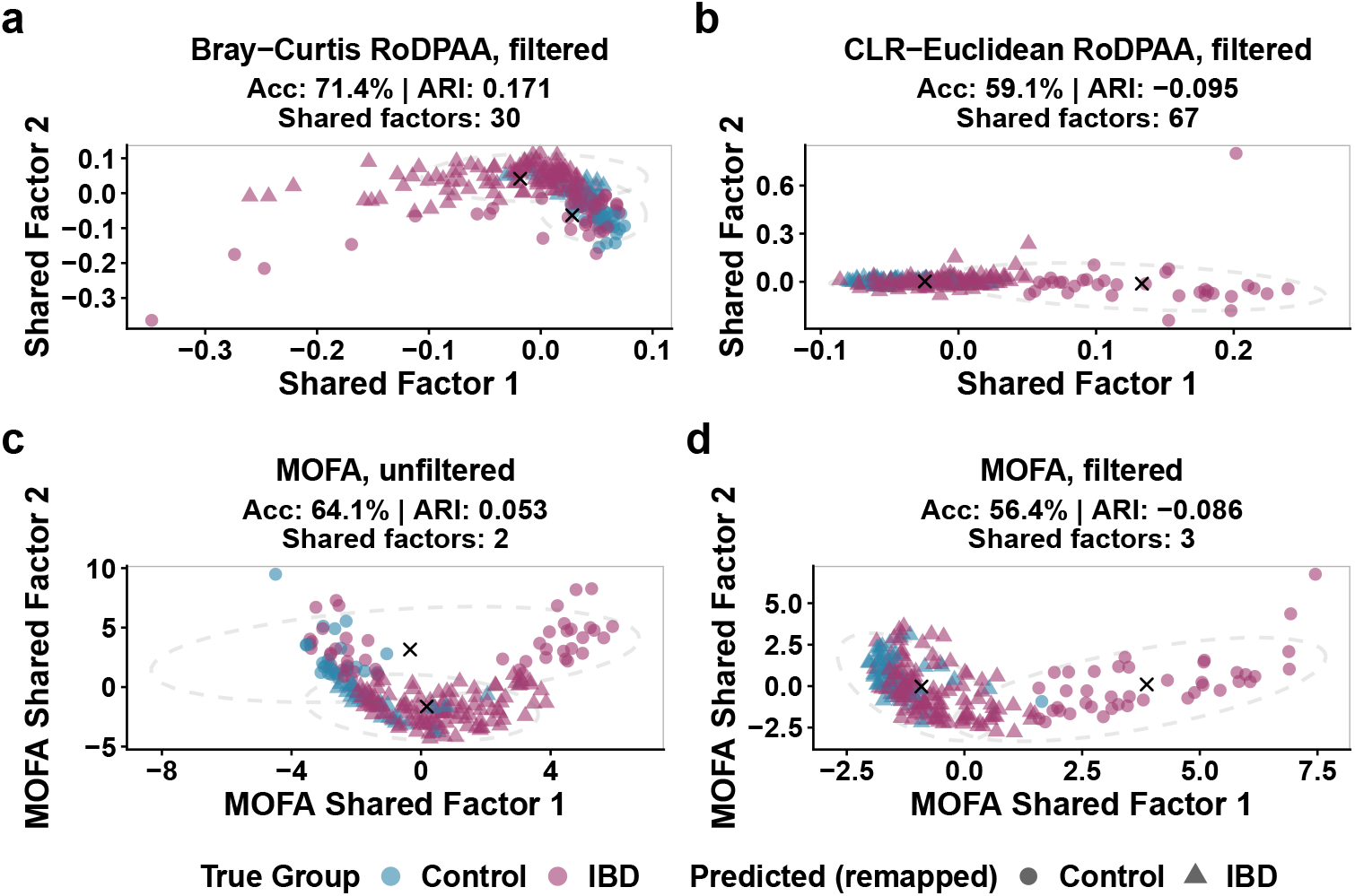
Sensitivity of RoDPAA and MOFA shared-factor recovery to feature filtering in the Franzosa IBD dataset. In panels a, b and d, microbial and metabolomic features observed in more than 80% of participants were retained, corresponding to a zero proportion below 20% (1,200 of 11,720 microbial and 3,973 of 8,848 metabolomic features); panel c uses all features. **a**, First two shared factors obtained using Bray–Curtis dis-similarity for the microbiome view and Euclidean distance for the log-transformed metabolomic view. RoDPAA selected a shared rank of 30, the same as without filtering. Unsupervised *k*-means clustering based on the first two factors achieved an accuracy of 71.4% and an adjusted Rand index (ARI) of 0.171 (76.4% and 0.267 without filtering; main-text Fig. 4a). **b**, First two shared factors obtained using Euclidean distance for both the CLR-transformed microbiome view and the log-transformed metabolomic view. RoDPAA selected a shared rank of 67 (0 without filtering), and clustering achieved an accuracy of 59.1% and an ARI of *−*0.095. **c**,**d**, First two MOFA shared factors without (**c**) and with (**d**) filtering. MOFA (MOFA2 default options, 15 initial factors) was fitted to the CLR-transformed microbiome and log-transformed metabolomic profiles; a factor was classified as shared when it explained more than 2% of the variance in both views, and the shared factors are ordered by total variance explained. MOFA identified two shared factors without filtering and three with filtering, and *k*-means clustering on the first two achieved an accuracy of 64.1% and an ARI of 0.053 without filtering, and 56.4% and *−*0.086 with filtering. The first two MOFA shared factors were similar with and without filtering (|*r*| = 0.90 and 0.80; Supplementary Table 2). Colors indicate the observed IBD and control groups, and point shapes indicate the estimated *k*-means clusters after permutation-invariant label matching; dashed ellipses are 95% data ellipses for the estimated clusters, and crosses mark cluster centers. Disease labels were not used in RoDPAA or MOFA estimation or *k*-means clustering.

**Extended Data Fig. 4.**
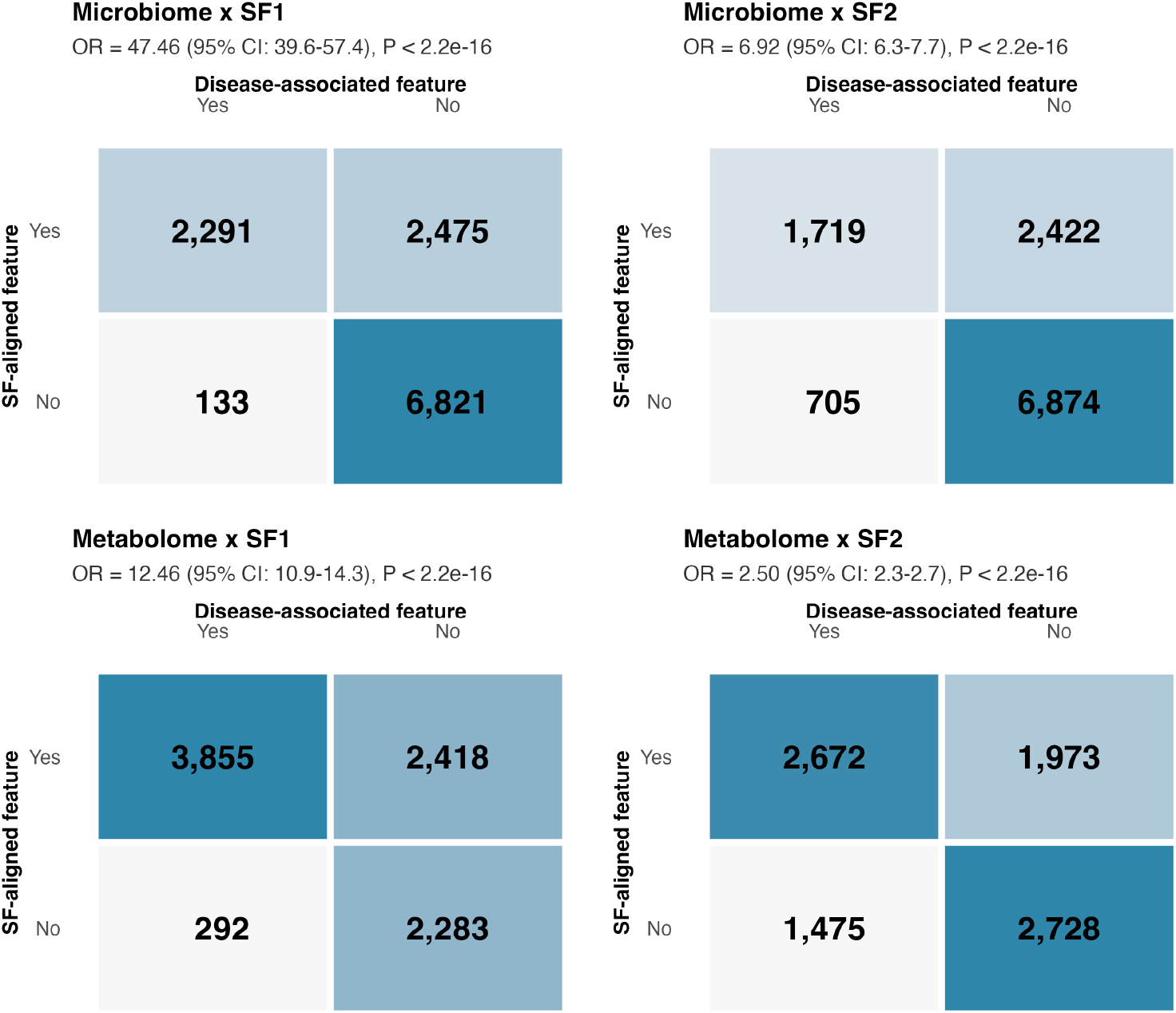
Feature-disease association versus feature-SF alignment (2*×*2 contingency tables). For each omics domain (microbiome, metabolome) and shared factor (SF1, SF2), features were classified as disease-associated (age-adjusted logistic regression, FDR *<* 0.05) and as SF-aligned (Spearman correlation with the shared-factor score, FDR *<* 0.05); microbiome and metabolome abundances were both log-transformed (log(*x* + pseudocount)) for these tests. Odds ratios (95% confidence intervals) were 47.46 (39.6–57.4) and 6.92 (6.3–7.7) for microbial features with SF1 and SF2, and 12.46 (10.9–14.3) and 2.50 (2.3–2.7) for metabolomic features (Fisher’s exact test, all *P <* 2.2 *×* 10^*−*16^). Inall four cases, features aligned with a shared factor were substantially more likely to also be disease-associated than non-aligned features.

**Extended Data Fig. 5.**
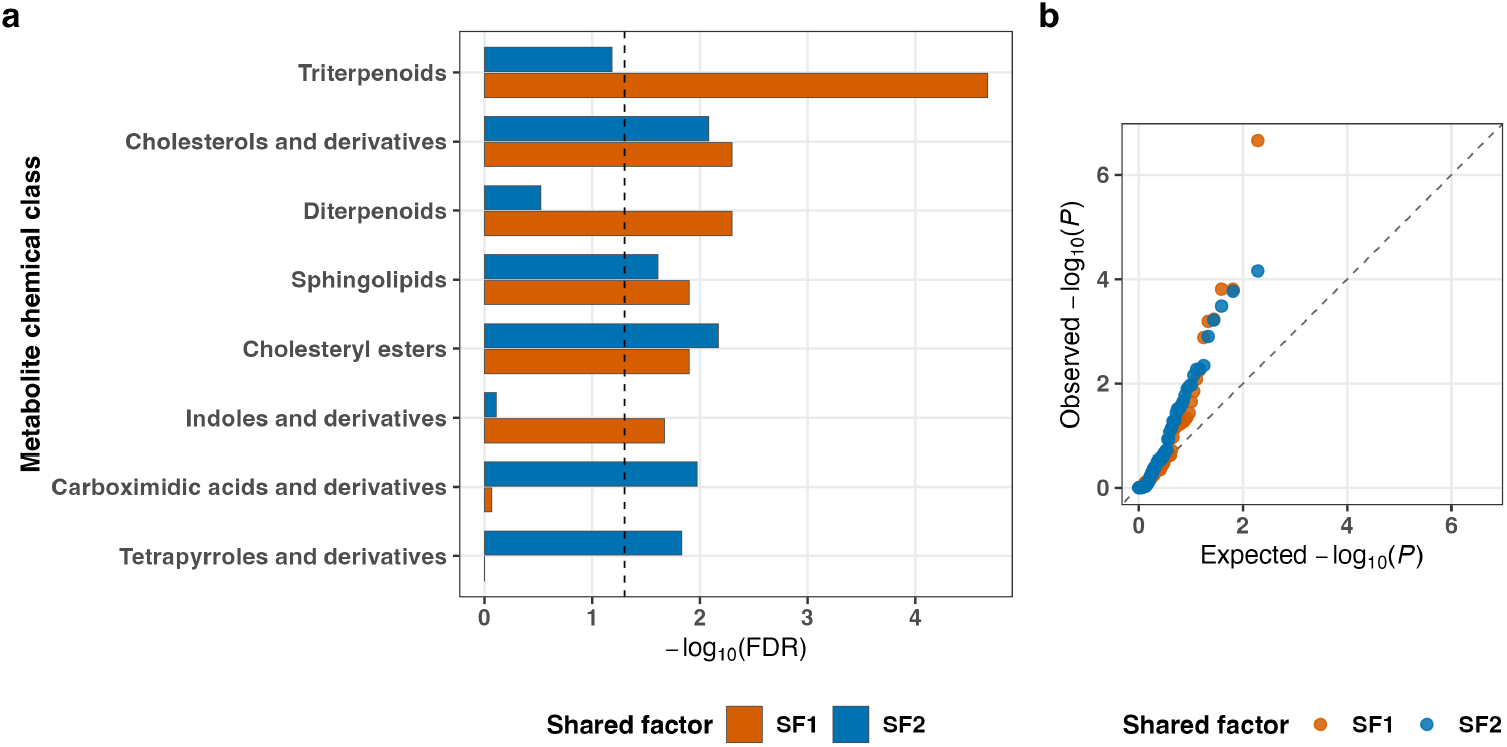
Metabolite chemical-class enrichment among SF_1_- and SF_2_-associated metabolites. **a**, For each putative chemical class (minimum 10 annotated metabolites; 97 classes tested), a hypergeometric test assessed over-representation among metabolites significantly associated (FDR *<* 0.05) with shared factor 1 (orange) or shared factor 2 (blue), out of all 3,820 annotated metabolites. Bars show *−* log_10_(FDR) for each class and factor; the dashed vertical line marks FDR = 0.05. Only classes significant in at least one factor are shown. **b**, Quantile-quantile plot of observed versus expected *−* log_10_(*P*) across all 97 tested classes for each factor, under the uniform-distribution null; the dashed diagonal line indicates the null expectation. Points departing above the diagonal correspond to the enriched classes shown in panel a. Metabolite significance calls use microbial and metabolomic features log-transformed identically, with false-discovery-rate correction pooled across both domains. Full results for all 97 classes are given in Supplementary Table 3.

## Supplementary Note 1: Euclidean Equivalence of RoDPAA and JIVE Signal Subspaces

Let **I**_*n*_ denote the *n× n* identity matrix, let **1***∈* ℝ^*n*^ denote the vector of ones, and define the centering matrix

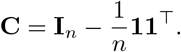

### Proposition 4.5.1

(Euclidean equivalence of RoDPAA and JIVE signal subspaces).

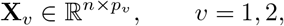

*denote two data views measured on the same n samples, and let* 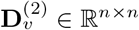 *denote the corresponding squared Euclidean distance matrices, with entries*

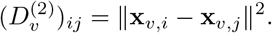

*Define the Gower-centered Euclidean similarity matrix*

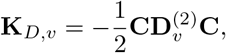

*and the centered data matrix*

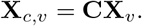

*Then, for each v* = 1, 2,

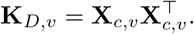

*Consequently, the signal eigenspace extracted by RoDPAA from* **K**_*D,v*_ *coincides with the corresponding left singular subspace of the centered data matrix* **X**_*c,v*_. *Therefore, under squared Euclidean distance, RoDPAA and the corresponding JIVE representation operate on the same sample-space signal subspaces and induce the same principal-angle geometry between views*.

*Proof* Fix a view *v ∈ {*1, 2*}*. For any two samples 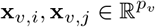, their squared Euclidean distance satisfies

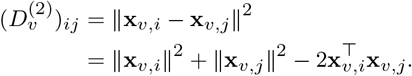

The matrix

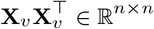

is the uncentered sample–sample Gram matrix associated with the linear kernel. Its (*i, j*) entry is

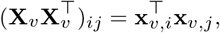

so 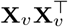 contains all pairwise inner-product similarities among the samples in view *v*. In particular,

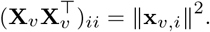

Define the vector of squared sample norms

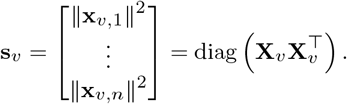

The squared Euclidean distance matrix can therefore be written as

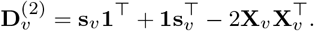

Applying Gower double-centering yields

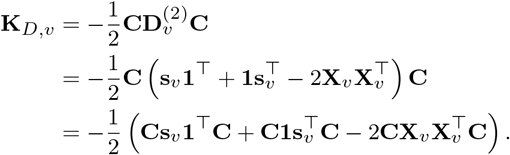

By the definition of the centering matrix,

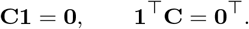

Hence,

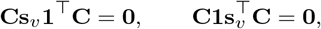

and therefore

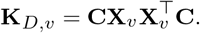

Thus, Gower double-centering removes the row- and column-specific squared-norm terms in the Euclidean distance matrix and retains the centered sample–sample inner-product structure.

Since

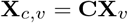

and **C**^*⊤*^ = **C**, we obtain

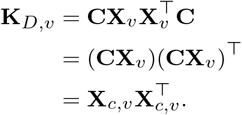

Therefore, under squared Euclidean distance, the centered similarity matrix used by RoDPAA is exactly the centered sample–sample Gram matrix, equivalently the centered linear kernel matrix. Its (*i, j*) entry is

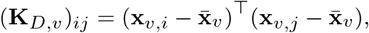

where

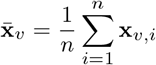

is the sample mean vector for view *v*.

Now let the singular value decomposition of the centered data matrix be

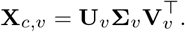

Then

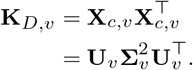

Hence, the eigenvectors of **K**_*D,v*_ associated with its positive eigenvalues are the left singular vectors of **X**_*c,v*_, and the corresponding eigenvalues are the squared singular values of **X**_*c,v*_.

Suppose that the same signal rank *r*_*v*_ is retained in the two representations. Let

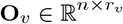

denote the orthonormal basis for the signal subspace extracted by RoDPAA from **K**_*D,v*_, and let

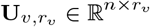

contain the corresponding leading *r*_*v*_ left singular vectors of **X**_*c,v*_. Then

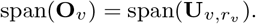

The particular orthonormal bases used to represent this common subspace need not be identical. Since **O**_*v*_ and 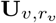 are orthonormal bases for the same *r*_*v*_-dimensional subspace, there exists an orthogonal matrix

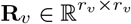

such that

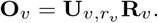

Applying this result to both views gives

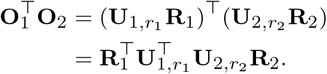

Because left and right multiplication by orthogonal matrices preserves singular values,

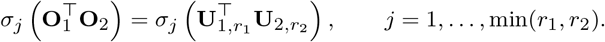

Therefore, under squared Euclidean distance and the same retained signal ranks, RoDPAA operates on the same sample-space signal subspaces as those obtained from the centered data matrices and induces the same principal-angle geometry between the two views.

Thus, the Euclidean signal-subspace construction underlying classical JIVE is embedded within the RoDPAA framework as a special case. RoDPAA extends this construction by allowing signal subspaces to be derived from centered similarity matrices constructed from user-specified pairwise dissimilarities.

## Supplementary Figures

**Supplementary Fig. 1.**
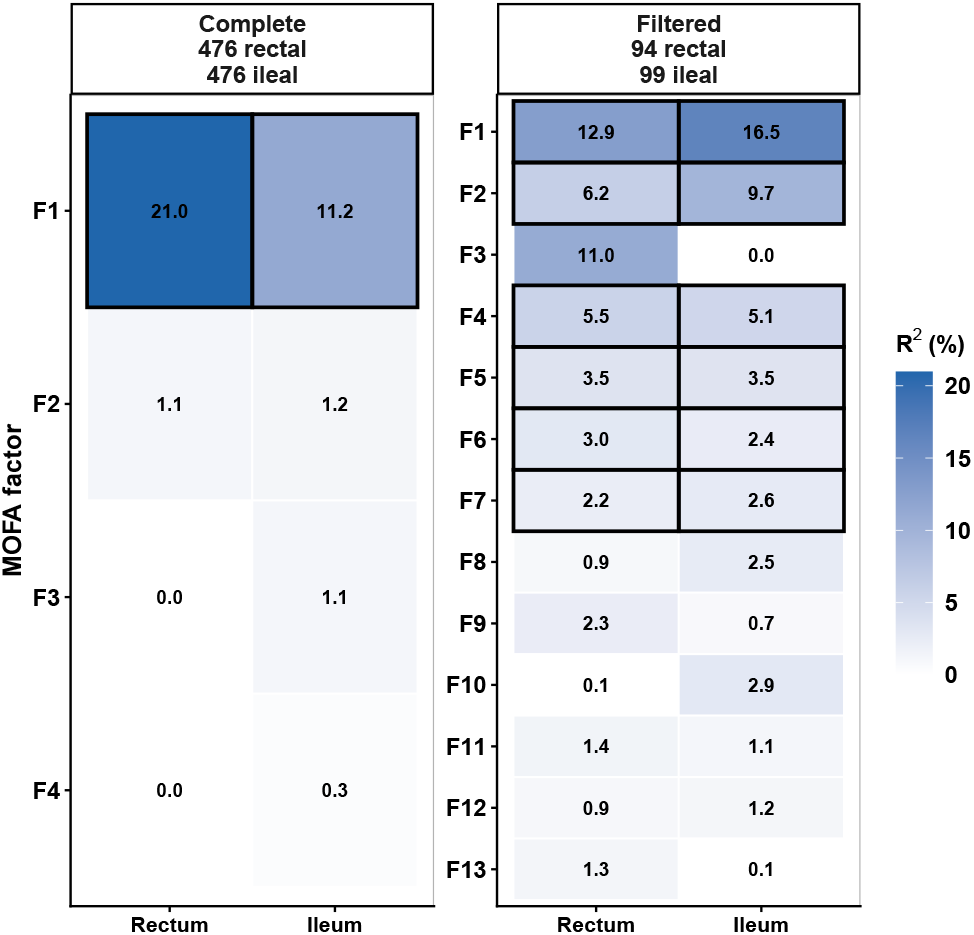
MOFA shared factors before and after prevalence filtering in the IBDMDB cohort. MOFA (`MOFA2` default options) was fitted to CLR-transformed rectal and ileal genus profiles of 68 participants. Variance explained by each retained factor in the complete data (476 rectal and 476 ileal genera) and after prevalence filtering (94 rectal and 99 ileal genera detected in at least 20% of participants). Outlined cells mark shared factors, defined as factors explaining more than 2% of the variance in both views: factor 1 in the complete data and factors 1, 2 and 4–7 after filtering.

**Supplementary Fig. 2.**
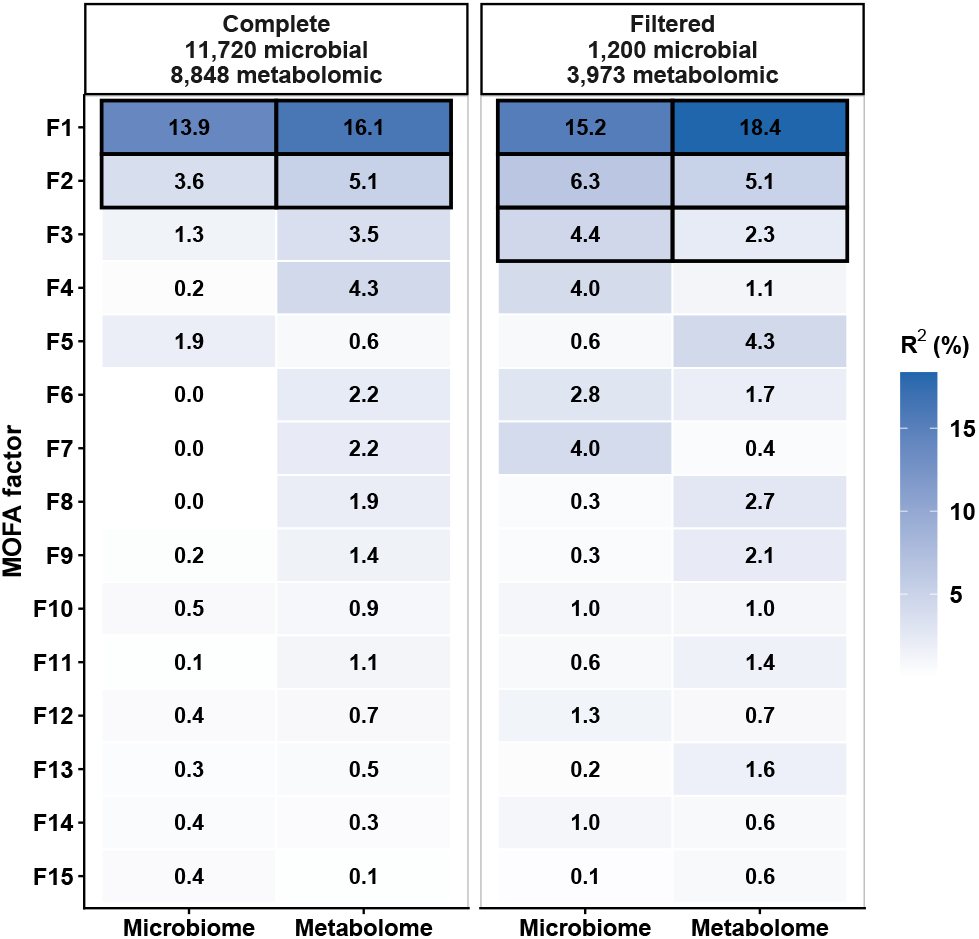
MOFA shared factors before and after prevalence filtering in the Franzosa cohort. Variance explained by each MOFA factor in the microbiome and metabolome views of 220 participants, in the complete data (11,720 microbial and 8,848 metabolomic features) and after the prevalence filtering of Extended Data Fig. 3 (features with a zero proportion below 20% in each view, 1,200 microbial and 3,973 metabolomic features). MOFA (`MOFA2` default options, 15 initial factors) was fitted to CLR-transformed microbiome and log-transformed metabolomic profiles. Outlined cells mark shared factors, defined as factors explaining more than 2% of the variance in both views: factors 1 and 2 in the complete data and factors 1–3 after filtering.

## Supplementary Algorithm

### Supplementary Algorithm 1

RoDPAA for two data views

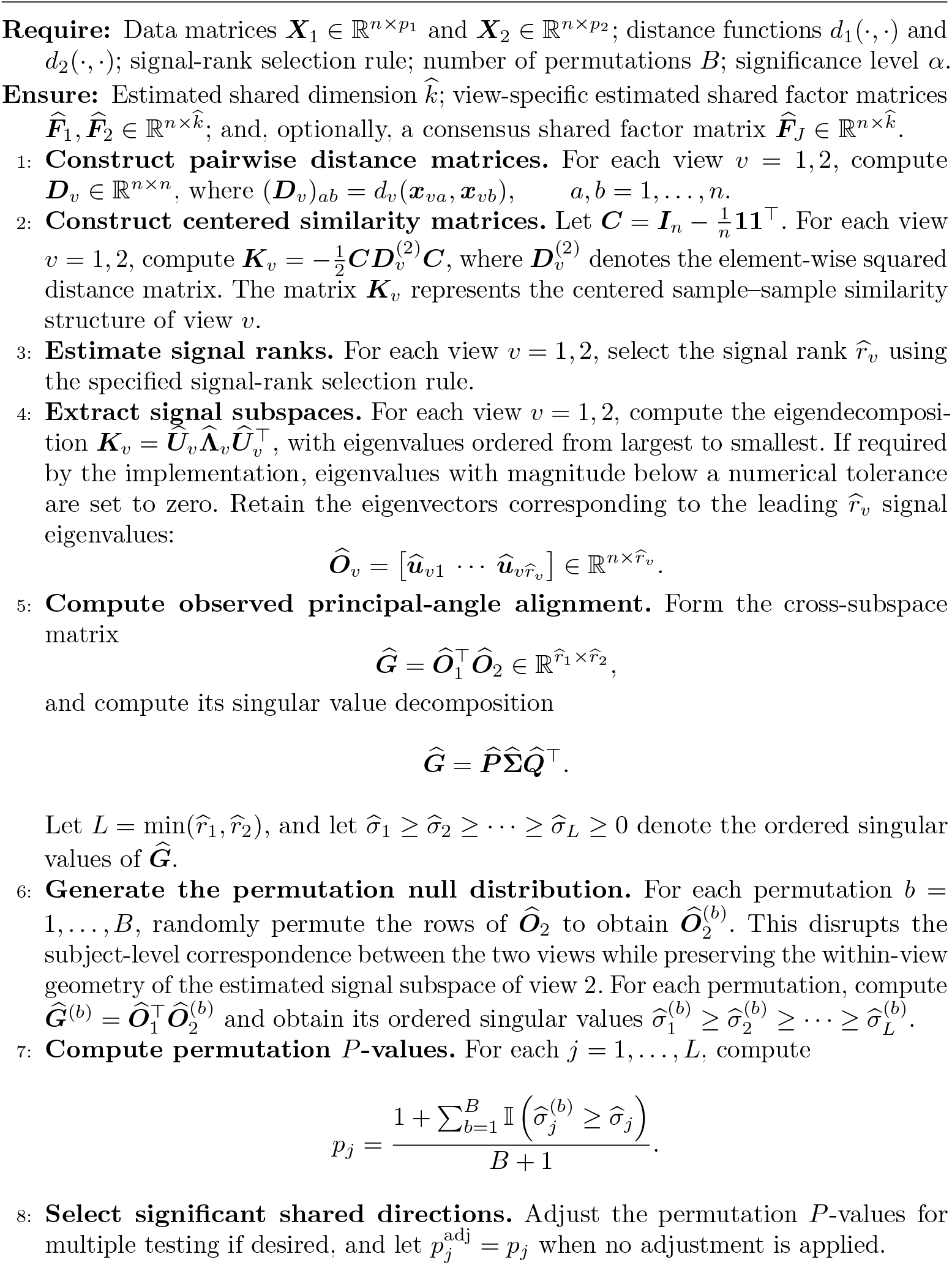
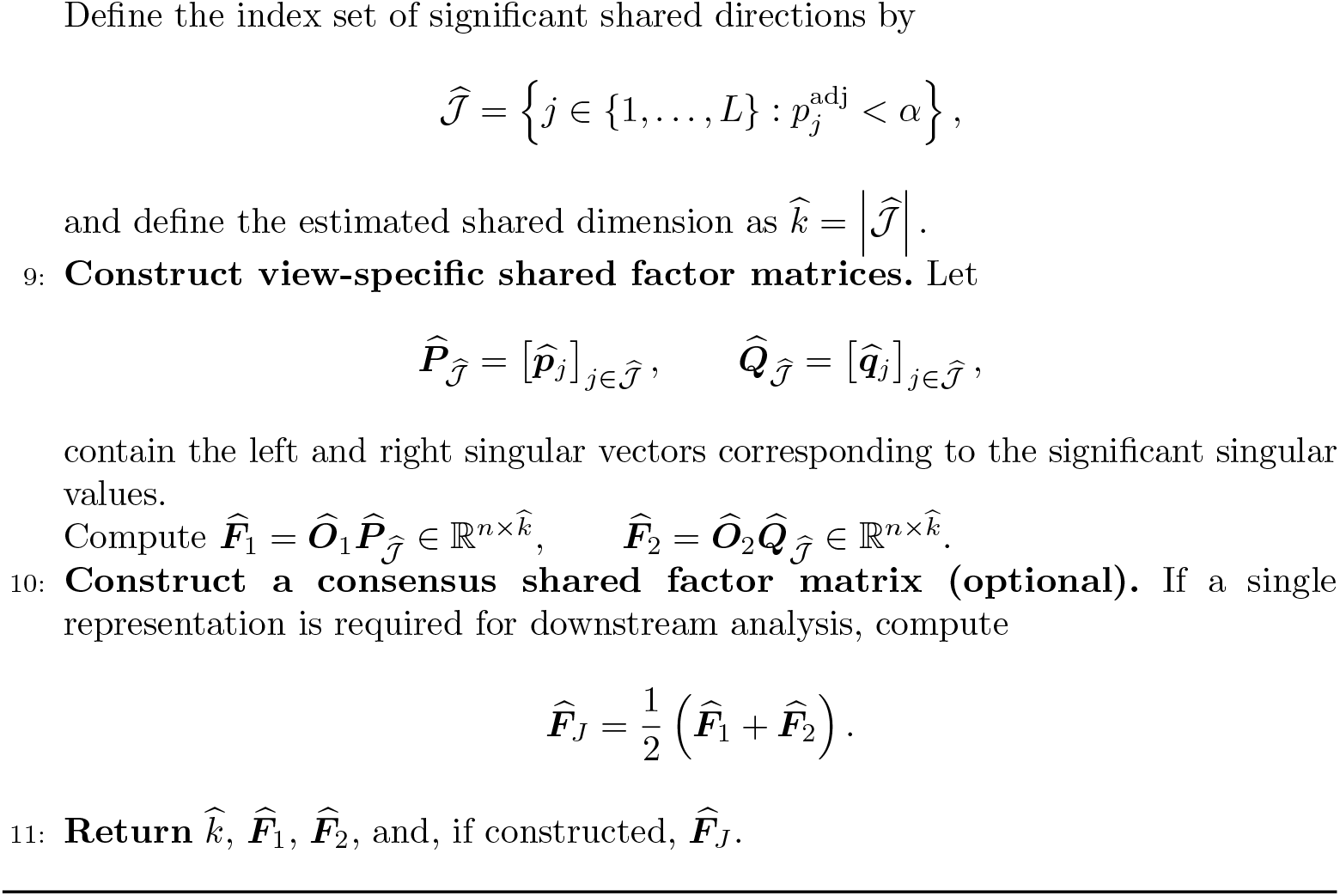

## Supplementary Tables

**Supplementary Table 1.**
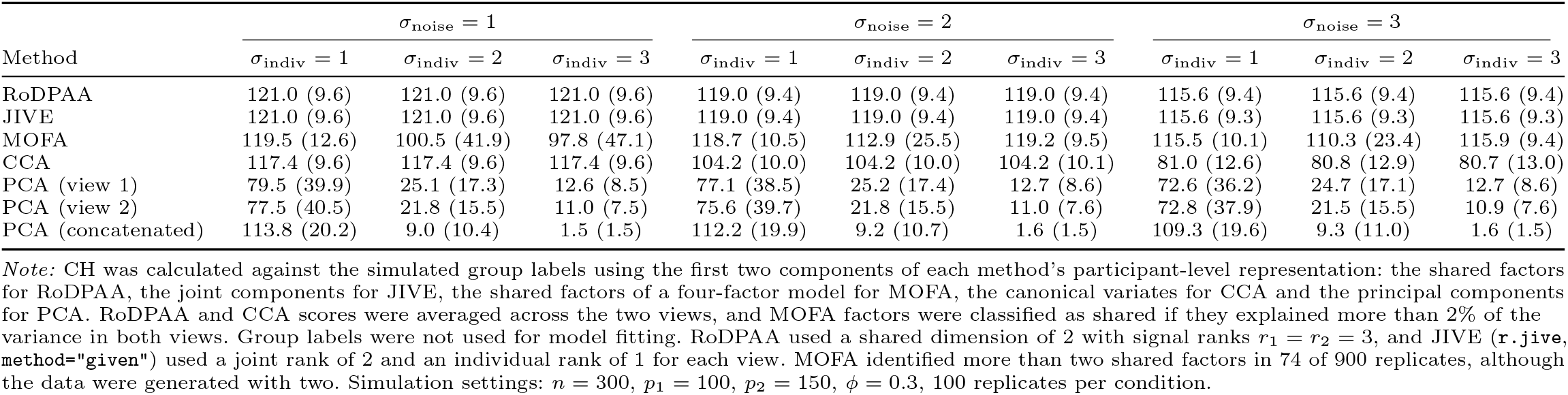
Calinski–Harabasz index across noise and view-specific variation in Simulation I. Mean (SD) Calinski–Harabasz (CH) index over 100 replicates for each combination of measurement noise (*σ*_noise_ *∈ {*1, 2, 3*}*) and view-specific variation (*σ*_indiv_ *∈ {*1, 2, 3*}*). The target dimension was prespecified as two, the dimension of the group-associated shared structure. Higher CH indicates better separation of the two simulated groups.

| Method | σ<sub>noise</sub> = 1 |  |  | σ<sub>noise</sub> = 2 |  |  | σ<sub>noise</sub> = 3 |  |  |
|---|---|---|---|---|---|---|---|---|---|
|  | σ<sub>indiv</sub> = 1 | σ<sub>indiv</sub> = 2 | σ<sub>indiv</sub> = 3 | σ<sub>indiv</sub> = 1 | σ<sub>indiv</sub> = 2 | σ<sub>indiv</sub> = 3 | σ<sub>indiv</sub> = 1 | σ<sub>indiv</sub> = 2 | σ<sub>indiv</sub> = 3 |
| RoDPAA | 121.0 (9.6) | 121.0 (9.6) | 121.0 (9.6) | 119.0 (9.4) | 119.0 (9.4) | 119.0 (9.4) | 115.6 (9.4) | 115.6 (9.4) | 115.6 (9.4) |
| JIVE | 121.0 (9.6) | 121.0 (9.6) | 121.0 (9.6) | 119.0 (9.4) | 119.0 (9.4) | 119.0 (9.4) | 115.6 (9.3) | 115.6 (9.3) | 115.6 (9.3) |
| MOFA | 119.5 (12.6) | 100.5 (41.9) | 97.8 (47.1) | 118.7 (10.5) | 112.9 (25.5) | 119.2 (9.5) | 115.5 (10.1) | 110.3 (23.4) | 115.9 (9.4) |
| CCA | 117.4 (9.6) | 117.4 (9.6) | 117.4 (9.6) | 104.2 (10.0) | 104.2 (10.0) | 104.2 (10.1) | 81.0 (12.6) | 80.8 (12.9) | 80.7 (13.0) |
| PCA (view 1) | 79.5 (39.9) | 25.1 (17.3) | 12.6 (8.5) | 77.1 (38.5) | 25.2 (17.4) | 12.7 (8.6) | 72.6 (36.2) | 24.7 (17.1) | 12.7 (8.6) |
| PCA (view 2) | 77.5 (40.5) | 21.8 (15.5) | 11.0 (7.5) | 75.6 (39.7) | 21.8 (15.5) | 11.0 (7.6) | 72.8 (37.9) | 21.5 (15.5) | 10.9 (7.6) |
| PCA (concatenated) | 113.8 (20.2) | 9.0 (10.4) | 1.5 (1.5) | 112.2 (19.9) | 9.2 (10.7) | 1.6 (1.5) | 109.3 (19.6) | 9.3 (11.0) | 1.6 (1.5) |

*Note:* CH was calculated against the simulated group labels using the first two components of each method's participant-level representation: the shared factors for RoDPAA, the joint components for JIVE, the shared factors of a four-factor model for MOFA, the canonical variates for CCA and the principal components for PCA. RoDPAA and CCA scores were averaged across the two views, and MOFA factors were classified as shared if they explained more than 2% of the variance in both views. Group labels were not used for model fitting. RoDPAA used a shared dimension of 2 with signal ranks *r*<sub>1</sub> = *r*<sub>2</sub> = 3, and JIVE (r.jive, method="given") used a joint rank of 2 and an individual rank of 1 for each view. MOFA identified more than two shared factors in 74 of 900 replicates, although the data were generated with two. Simulation settings: *n* = 300, *p*<sub>1</sub> = 100, *p*<sub>2</sub> = 150, *ϕ* = 0.3, 100 replicates per condition.

**Supplementary Table 2.**
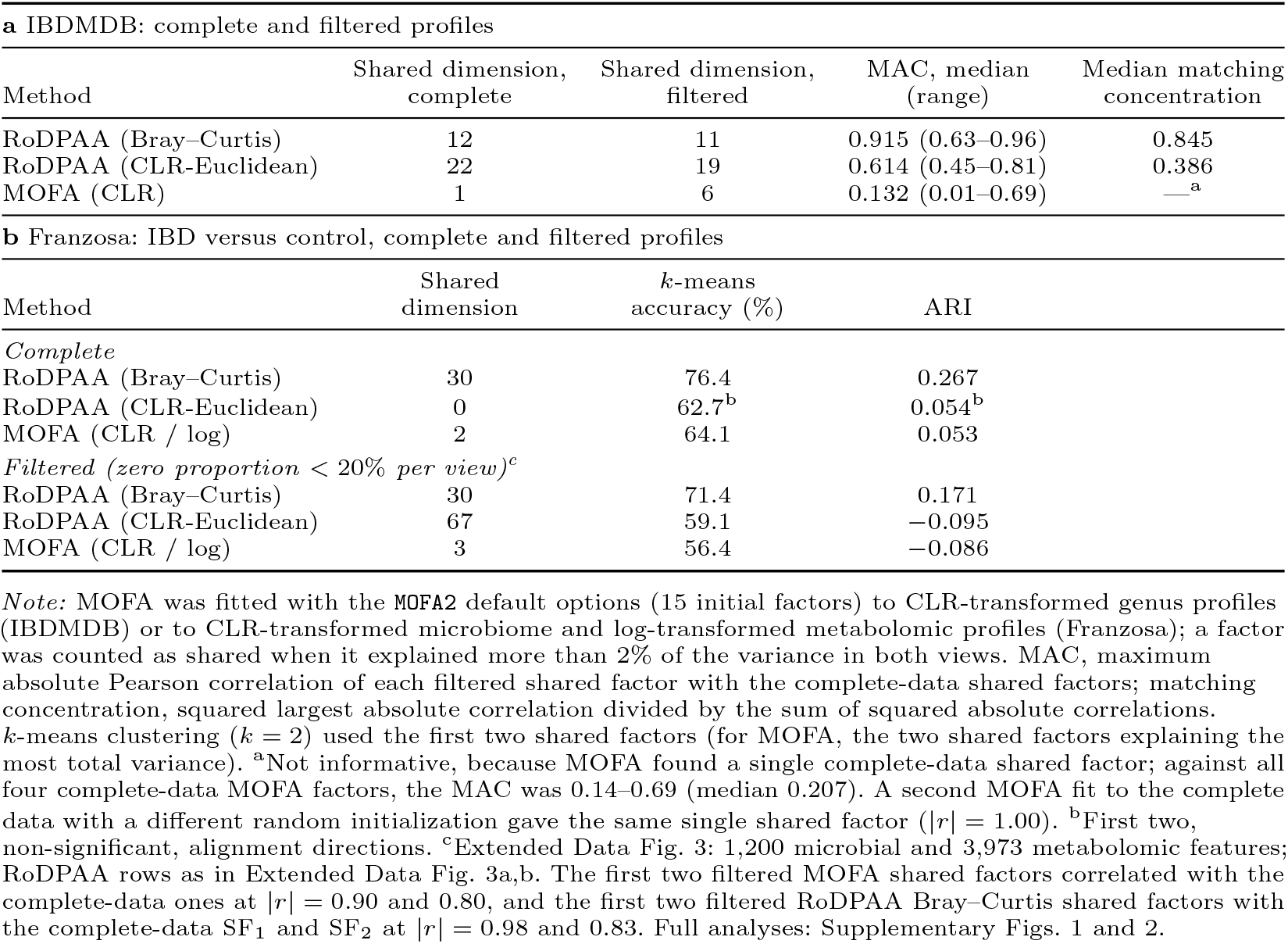
RoDPAA and MOFA in the two IBD applications. **a**, Stability of the shared structure to prevalence filtering in the IBDMDB cohort (ileum and rectum genus profiles, 68 participants). **b**, Separation of IBD and control participants in the Franzosa cohort (microbiome and metabolome, 220 participants), without and with the prevalence filtering of Extended Data Fig. 3.

| **a** IBDMDB: complete and filtered profiles |  |  |  |  |
|---|---|---|---|---|
| Method | Shared dimension, complete | Shared dimension, filtered | MAC, median (range) | Median matching concentration |
| RoDPAA (Bray–Curtis) | 12 | 11 | 0.915 (0.63–0.96) | 0.845 |
| RoDPAA (CLR-Euclidean) | 22 | 19 | 0.614 (0.45–0.81) | 0.386 |
| MOFA (CLR) | 1 | 6 | 0.132 (0.01–0.69) | —<sup>a</sup> |
| **b** Franzosa: IBD versus control, complete and filtered profiles |  |  |  |  |
| Method | Shared dimension | *k*-means accuracy (%) | ARI |  |
| *Complete* |  |  |  |  |
| RoDPAA (Bray–Curtis) | 30 | 76.4 | 0.267 |  |
| RoDPAA (CLR-Euclidean) | 0 | 62.7<sup>b</sup> | 0.054<sup>b</sup> |  |
| MOFA (CLR / log) | 2 | 64.1 | 0.053 |  |
| *Filtered (zero proportion < 20% per view)*<sup>c</sup> |  |  |  |  |
| RoDPAA (Bray–Curtis) | 30 | 71.4 | 0.171 |  |
| RoDPAA (CLR-Euclidean) | 67 | 59.1 | −0.095 |  |
| MOFA (CLR / log) | 3 | 56.4 | −0.086 |  |

*Note:* MOFA was fitted with the MOFA2 default options (15 initial factors) to CLR-transformed genus profiles (IBDMDB) or to CLR-transformed microbiome and log-transformed metabolomic profiles (Franzosa); a factor was counted as shared when it explained more than 2% of the variance in both views. MAC, maximum absolute Pearson correlation of each filtered shared factor with the complete-data shared factors; matching concentration, squared largest absolute correlation divided by the sum of squared absolute correlations. *k*-means clustering (*k* = 2) used the first two shared factors (for MOFA, the two shared factors explaining the most total variance). <sup>a</sup>Not informative, because MOFA found a single complete-data shared factor; against all four complete-data MOFA factors, the MAC was 0.14–0.69 (median 0.207). A second MOFA fit to the complete data with a different random initialization gave the same single shared factor (\|*r*\| = 1.00). <sup>b</sup>First two, non-significant, alignment directions. <sup>c</sup>Extended Data Fig. 3: 1,200 microbial and 3,973 metabolomic features; RoDPAA rows as in Extended Data Fig. 3a,b. The first two filtered MOFA shared factors correlated with the complete-data ones at \|*r*\| = 0.90 and 0.80, and the first two filtered RoDPAA Bray–Curtis shared factors with the complete-data SF<sub>1</sub> and SF<sub>2</sub> at \|*r*\| = 0.98 and 0.83. Full analyses: Supplementary Figs. 1 and 2.

**Supplementary Table 3.**
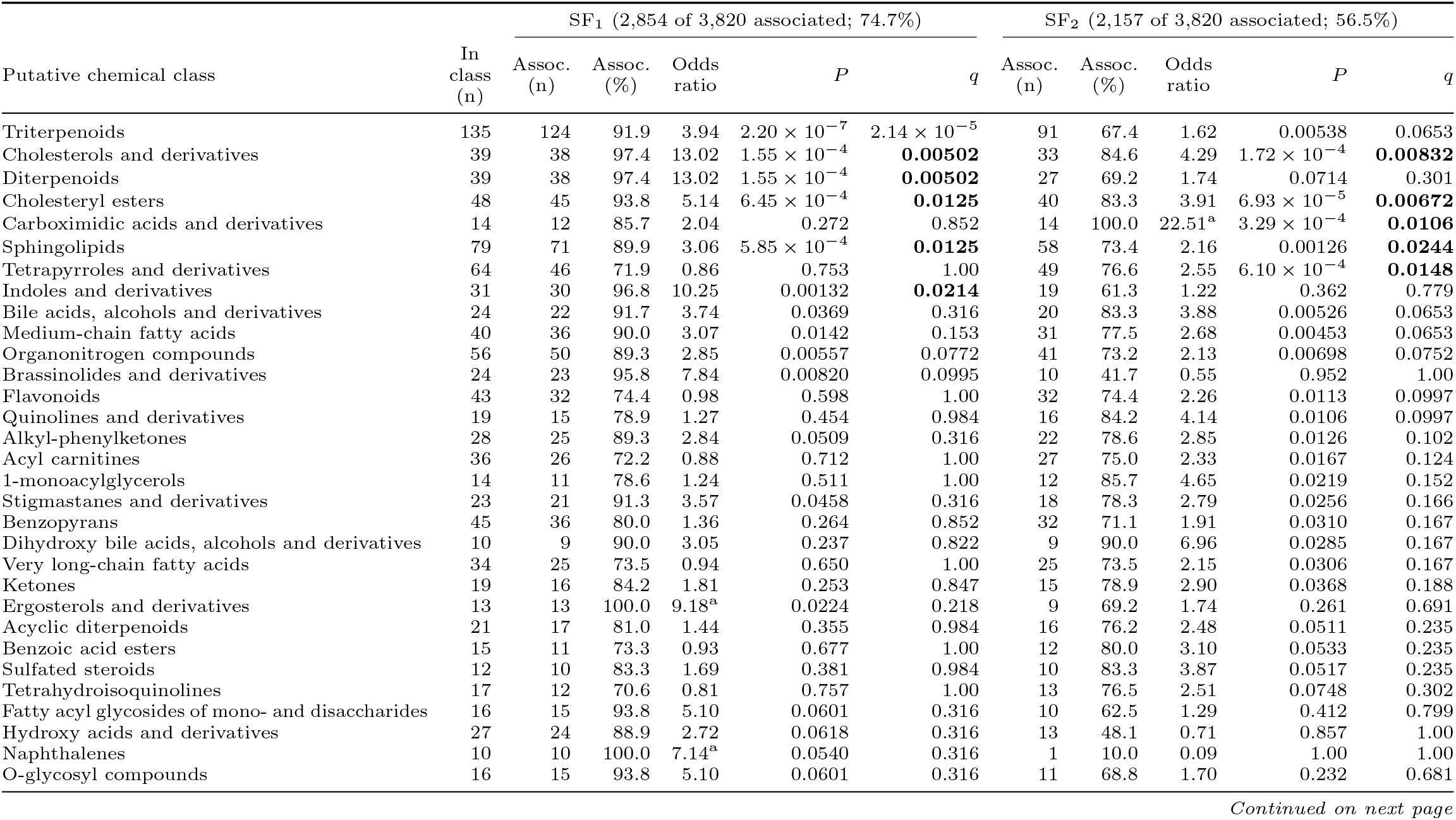
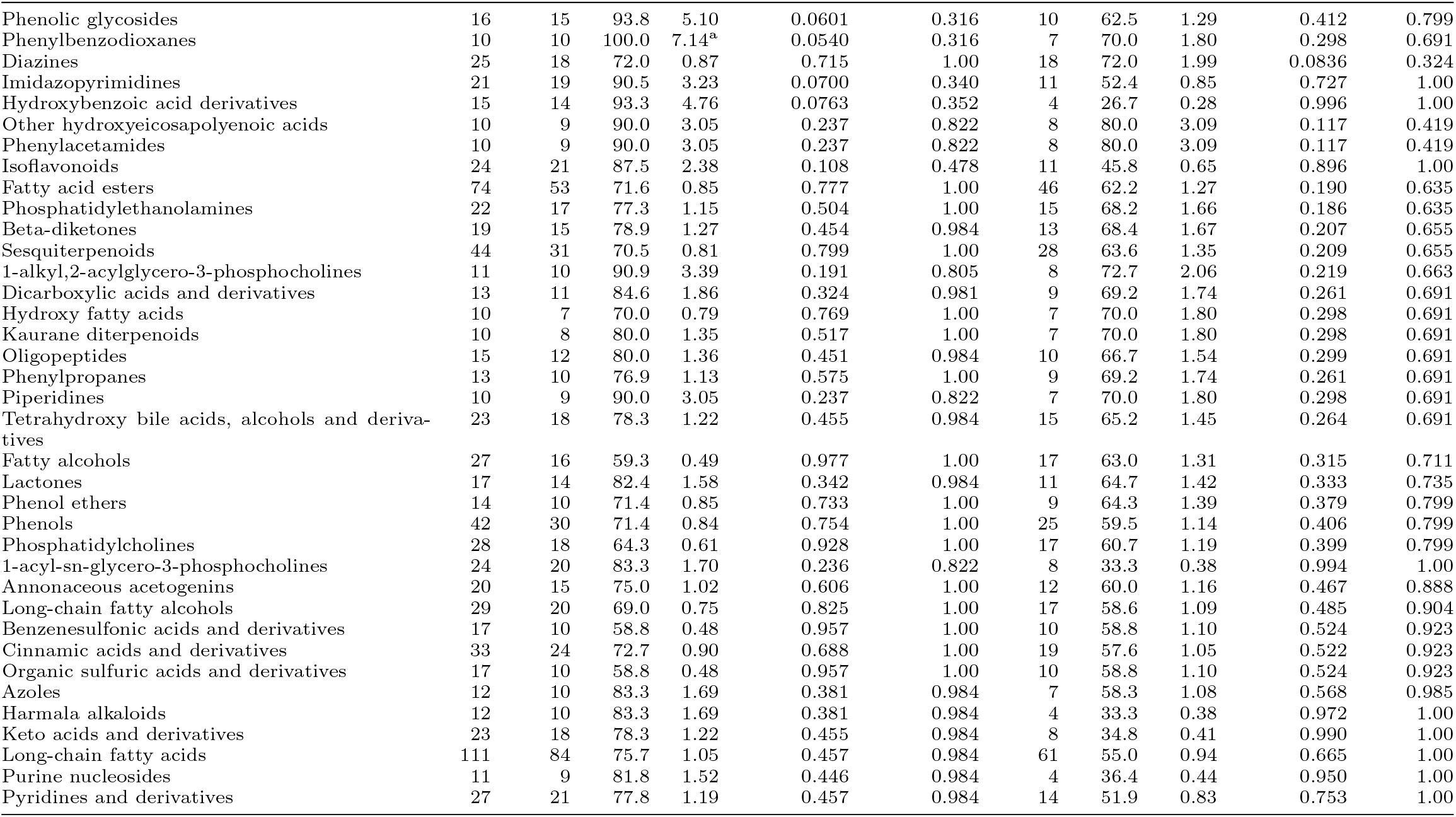
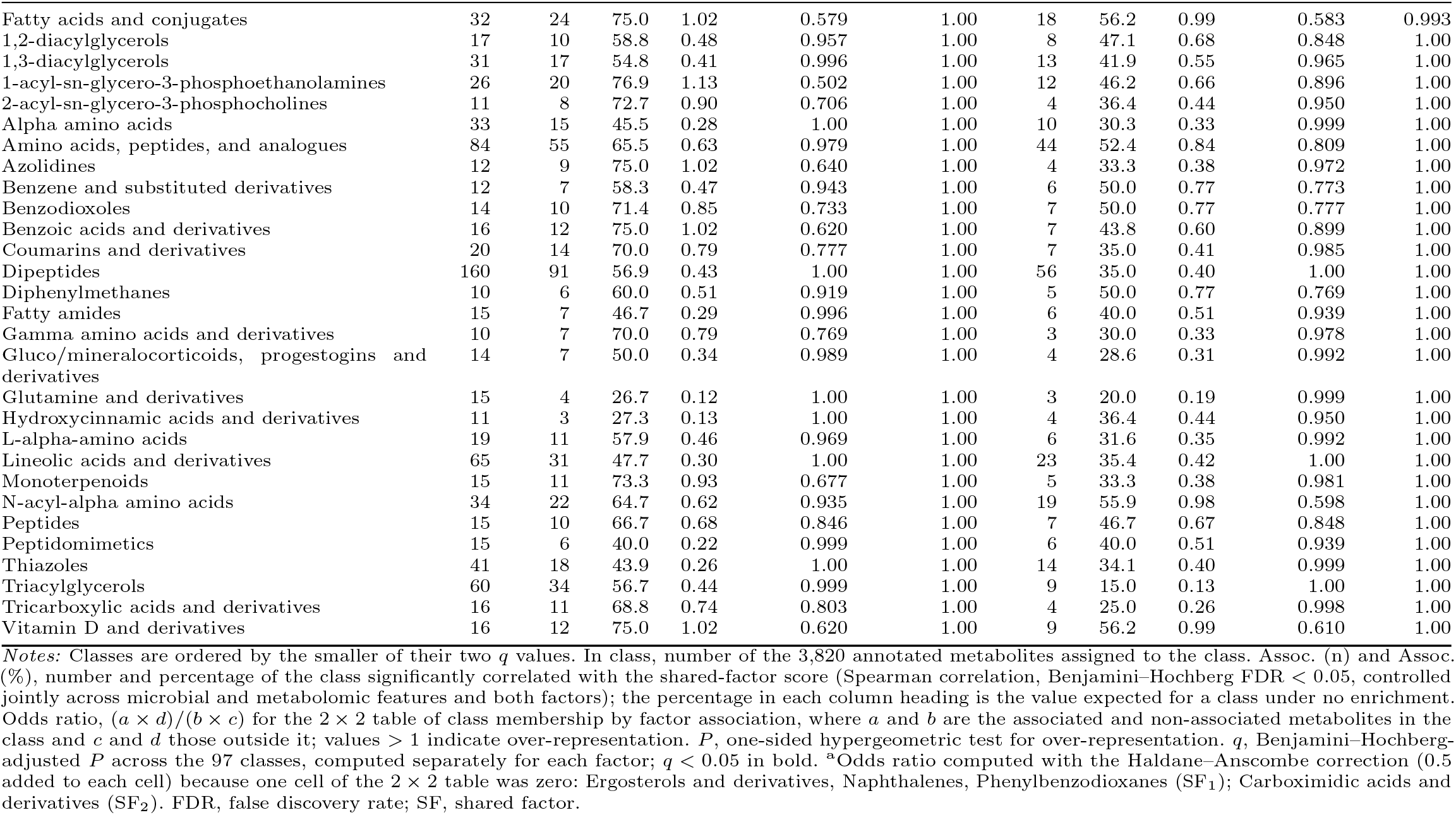
Chemical-class enrichment of metabolites associated with shared factors SF_1_ and SF_2_ in the Franzosa IBD cohort. For each of the 97 putative chemical classes with *≥*10 annotated metabolites, we tested whether the class contained more metabolites significantly correlated with each RoDPAA shared factor (Spearman correlation, FDR *<* 0.05) than expected by chance, using all 3,820 annotated metabolites as background (2,854 associated with SF_1_ and 2,157 with SF_2_). These results underlie Extended Data Fig. 5. The same table is provided as a separate Excel file (Supplementary Table 3 metabolite class enrichment.xlsx).

| Putative chemical class | In class (n) | SF<sub>1</sub> (2,854 of 3,820 associated; 74.7%) |  |  |  |  | SF<sub>2</sub> (2,157 of 3,820 associated; 56.5%) |  |  |  |  |
|---|---|---|---|---|---|---|---|---|---|---|---|
|  |  | Assoc. (n) | Assoc. (%) | Odds ratio | *P* | *q* | Assoc. (n) | Assoc. (%) | Odds ratio | *P* | *q* |
| Triterpenoids | 135 | 124 | 91.9 | 3.94 | 2.20 × 10<sup>−7</sup> | 2.14 × 10<sup>−5</sup> | 91 | 67.4 | 1.62 | 0.00538 | 0.0653 |
| Cholesterols and derivatives | 39 | 38 | 97.4 | 13.02 | 1.55 × 10<sup>−4</sup> | **0.00502** | 33 | 84.6 | 4.29 | 1.72 × 10<sup>−4</sup> | **0.00832** |
| Diterpenoids | 39 | 38 | 97.4 | 13.02 | 1.55 × 10<sup>−4</sup> | **0.00502** | 27 | 69.2 | 1.74 | 0.0714 | 0.301 |
| Cholesteryl esters | 48 | 45 | 93.8 | 5.14 | 6.45 × 10<sup>−4</sup> | **0.0125** | 40 | 83.3 | 3.91 | 6.93 × 10<sup>−5</sup> | **0.00672** |
| Carboximidic acids and derivatives | 14 | 12 | 85.7 | 2.04 | 0.272 | 0.852 | 14 | 100.0 | 22.51<sup>a</sup> | 3.29 × 10<sup>−4</sup> | **0.0106** |
| Sphingolipids | 79 | 71 | 89.9 | 3.06 | 5.85 × 10<sup>−4</sup> | **0.0125** | 58 | 73.4 | 2.16 | 0.00126 | **0.0244** |
| Tetrapyrroles and derivatives | 64 | 46 | 71.9 | 0.86 | 0.753 | 1.00 | 49 | 76.6 | 2.55 | 6.10 × 10<sup>−4</sup> | **0.0148** |
| Indoles and derivatives | 31 | 30 | 96.8 | 10.25 | 0.00132 | **0.0214** | 19 | 61.3 | 1.22 | 0.362 | 0.779 |
| Bile acids, alcohols and derivatives | 24 | 22 | 91.7 | 3.74 | 0.0369 | 0.316 | 20 | 83.3 | 3.88 | 0.00526 | 0.0653 |
| Medium-chain fatty acids | 40 | 36 | 90.0 | 3.07 | 0.0142 | 0.153 | 31 | 77.5 | 2.68 | 0.00453 | 0.0653 |
| Organonitrogen compounds | 56 | 50 | 89.3 | 2.85 | 0.00557 | 0.0772 | 41 | 73.2 | 2.13 | 0.00698 | 0.0752 |
| Brassinolides and derivatives | 24 | 23 | 95.8 | 7.84 | 0.00820 | 0.0995 | 10 | 41.7 | 0.55 | 0.952 | 1.00 |
| Flavonoids | 43 | 32 | 74.4 | 0.98 | 0.598 | 1.00 | 32 | 74.4 | 2.26 | 0.0113 | 0.0997 |
| Quinolines and derivatives | 19 | 15 | 78.9 | 1.27 | 0.454 | 0.984 | 16 | 84.2 | 4.14 | 0.0106 | 0.0997 |
| Alkyl-phenylketones | 28 | 25 | 89.3 | 2.84 | 0.0509 | 0.316 | 22 | 78.6 | 2.85 | 0.0126 | 0.102 |
| Acyl carnitines | 36 | 26 | 72.2 | 0.88 | 0.712 | 1.00 | 27 | 75.0 | 2.33 | 0.0167 | 0.124 |
| 1-monoacylglycerols | 14 | 11 | 78.6 | 1.24 | 0.511 | 1.00 | 12 | 85.7 | 4.65 | 0.0219 | 0.152 |
| Stigmastanes and derivatives | 23 | 21 | 91.3 | 3.57 | 0.0458 | 0.316 | 18 | 78.3 | 2.79 | 0.0256 | 0.166 |
| Benzopyrans | 45 | 36 | 80.0 | 1.36 | 0.264 | 0.852 | 32 | 71.1 | 1.91 | 0.0310 | 0.167 |
| Dihydroxy bile acids, alcohols and derivatives | 10 | 9 | 90.0 | 3.05 | 0.237 | 0.822 | 9 | 90.0 | 6.96 | 0.0285 | 0.167 |
| Very long-chain fatty acids | 34 | 25 | 73.5 | 0.94 | 0.650 | 1.00 | 25 | 73.5 | 2.15 | 0.0306 | 0.167 |
| Ketones | 19 | 16 | 84.2 | 1.81 | 0.253 | 0.847 | 15 | 78.9 | 2.90 | 0.0368 | 0.188 |
| Ergosterols and derivatives | 13 | 13 | 100.0 | 9.18<sup>a</sup> | 0.0224 | 0.218 | 9 | 69.2 | 1.74 | 0.261 | 0.691 |
| Acyclic diterpenoids | 21 | 17 | 81.0 | 1.44 | 0.355 | 0.984 | 16 | 76.2 | 2.48 | 0.0511 | 0.235 |
| Benzoic acid esters | 15 | 11 | 73.3 | 0.93 | 0.677 | 1.00 | 12 | 80.0 | 3.10 | 0.0533 | 0.235 |
| Sulfated steroids | 12 | 10 | 83.3 | 1.69 | 0.381 | 0.984 | 10 | 83.3 | 3.87 | 0.0517 | 0.235 |
| Tetrahydroisoquinolines | 17 | 12 | 70.6 | 0.81 | 0.757 | 1.00 | 13 | 76.5 | 2.51 | 0.0748 | 0.302 |
| Fatty acyl glycosides of mono- and disaccharides | 16 | 15 | 93.8 | 5.10 | 0.0601 | 0.316 | 10 | 62.5 | 1.29 | 0.412 | 0.799 |
| Hydroxy acids and derivatives | 27 | 24 | 88.9 | 2.72 | 0.0618 | 0.316 | 13 | 48.1 | 0.71 | 0.857 | 1.00 |
| Naphthalenes | 10 | 10 | 100.0 | 7.14<sup>a</sup> | 0.0540 | 0.316 | 1 | 10.0 | 0.09 | 1.00 | 1.00 |
| O-glycosyl compounds | 16 | 15 | 93.8 | 5.10 | 0.0601 | 0.316 | 11 | 68.8 | 1.70 | 0.232 | 0.681 |
| Phenolic glycosides | 16 | 15 | 93.8 | 5.10 | 0.0601 | 0.316 | 10 | 62.5 | 1.29 | 0.412 | 0.799 |
| Phenylbenzodioxanes | 10 | 10 | 100.0 | 7.14<sup>a</sup> | 0.0540 | 0.316 | 7 | 70.0 | 1.80 | 0.298 | 0.691 |
| Diazines | 25 | 18 | 72.0 | 0.87 | 0.715 | 1.00 | 18 | 72.0 | 1.99 | 0.0836 | 0.324 |
| Imidazopyrimidines | 21 | 19 | 90.5 | 3.23 | 0.0700 | 0.340 | 11 | 52.4 | 0.85 | 0.727 | 1.00 |
| Hydroxybenzoic acid derivatives | 15 | 14 | 93.3 | 4.76 | 0.0763 | 0.352 | 4 | 26.7 | 0.28 | 0.996 | 1.00 |
| Other hydroxyeicosapolyenoic acids | 10 | 9 | 90.0 | 3.05 | 0.237 | 0.822 | 8 | 80.0 | 3.09 | 0.117 | 0.419 |
| Phenylacetamides | 10 | 9 | 90.0 | 3.05 | 0.237 | 0.822 | 8 | 80.0 | 3.09 | 0.117 | 0.419 |
| Isoflavonoids | 24 | 21 | 87.5 | 2.38 | 0.108 | 0.478 | 11 | 45.8 | 0.65 | 0.896 | 1.00 |
| Fatty acid esters | 74 | 53 | 71.6 | 0.85 | 0.777 | 1.00 | 46 | 62.2 | 1.27 | 0.190 | 0.635 |
| Phosphatidylethanolamines | 22 | 17 | 77.3 | 1.15 | 0.504 | 1.00 | 15 | 68.2 | 1.66 | 0.186 | 0.635 |
| Beta-diketones | 19 | 15 | 78.9 | 1.27 | 0.454 | 0.984 | 13 | 68.4 | 1.67 | 0.207 | 0.655 |
| Sesquiterpenoids | 44 | 31 | 70.5 | 0.81 | 0.799 | 1.00 | 28 | 63.6 | 1.35 | 0.209 | 0.655 |
| 1-alkyl,2-acylglycero-3-phosphocholines | 11 | 10 | 90.9 | 3.39 | 0.191 | 0.805 | 8 | 72.7 | 2.06 | 0.219 | 0.663 |
| Dicarboxylic acids and derivatives | 13 | 11 | 84.6 | 1.86 | 0.324 | 0.981 | 9 | 69.2 | 1.74 | 0.261 | 0.691 |
| Hydroxy fatty acids | 10 | 7 | 70.0 | 0.79 | 0.769 | 1.00 | 7 | 70.0 | 1.80 | 0.298 | 0.691 |
| Kaurane diterpenoids | 10 | 8 | 80.0 | 1.35 | 0.517 | 1.00 | 7 | 70.0 | 1.80 | 0.298 | 0.691 |
| Oligopeptides | 15 | 12 | 80.0 | 1.36 | 0.451 | 0.984 | 10 | 66.7 | 1.54 | 0.299 | 0.691 |
| Phenylpropanes | 13 | 10 | 76.9 | 1.13 | 0.575 | 1.00 | 9 | 69.2 | 1.74 | 0.261 | 0.691 |
| Piperidines | 10 | 9 | 90.0 | 3.05 | 0.237 | 0.822 | 7 | 70.0 | 1.80 | 0.298 | 0.691 |
| Tetrahydroxy bile acids, alcohols and deriva-tives | 23 | 18 | 78.3 | 1.22 | 0.455 | 0.984 | 15 | 65.2 | 1.45 | 0.264 | 0.691 |
| Fatty alcohols | 27 | 16 | 59.3 | 0.49 | 0.977 | 1.00 | 17 | 63.0 | 1.31 | 0.315 | 0.711 |
| Lactones | 17 | 14 | 82.4 | 1.58 | 0.342 | 0.984 | 11 | 64.7 | 1.42 | 0.333 | 0.735 |
| Phenol ethers | 14 | 10 | 71.4 | 0.85 | 0.733 | 1.00 | 9 | 64.3 | 1.39 | 0.379 | 0.799 |
| Phenols | 42 | 30 | 71.4 | 0.84 | 0.754 | 1.00 | 25 | 59.5 | 1.14 | 0.406 | 0.799 |
| Phosphatidylcholines | 28 | 18 | 64.3 | 0.61 | 0.928 | 1.00 | 17 | 60.7 | 1.19 | 0.399 | 0.799 |
| 1-acyl-sn-glycero-3-phosphocholines | 24 | 20 | 83.3 | 1.70 | 0.236 | 0.822 | 8 | 33.3 | 0.38 | 0.994 | 1.00 |
| Annonaceous acetogenins | 20 | 15 | 75.0 | 1.02 | 0.606 | 1.00 | 12 | 60.0 | 1.16 | 0.467 | 0.888 |
| Long-chain fatty alcohols | 29 | 20 | 69.0 | 0.75 | 0.825 | 1.00 | 17 | 58.6 | 1.09 | 0.485 | 0.904 |
| Benzenesulfonic acids and derivatives | 17 | 10 | 58.8 | 0.48 | 0.957 | 1.00 | 10 | 58.8 | 1.10 | 0.524 | 0.923 |
| Cinnamic acids and derivatives | 33 | 24 | 72.7 | 0.90 | 0.688 | 1.00 | 19 | 57.6 | 1.05 | 0.522 | 0.923 |
| Organic sulfuric acids and derivatives | 17 | 10 | 58.8 | 0.48 | 0.957 | 1.00 | 10 | 58.8 | 1.10 | 0.524 | 0.923 |
| Azoles | 12 | 10 | 83.3 | 1.69 | 0.381 | 0.984 | 7 | 58.3 | 1.08 | 0.568 | 0.985 |
| Harmala alkaloids | 12 | 10 | 83.3 | 1.69 | 0.381 | 0.984 | 4 | 33.3 | 0.38 | 0.972 | 1.00 |
| Keto acids and derivatives | 23 | 18 | 78.3 | 1.22 | 0.455 | 0.984 | 8 | 34.8 | 0.41 | 0.990 | 1.00 |
| Long-chain fatty acids | 111 | 84 | 75.7 | 1.05 | 0.457 | 0.984 | 61 | 55.0 | 0.94 | 0.665 | 1.00 |
| Purine nucleosides | 11 | 9 | 81.8 | 1.52 | 0.446 | 0.984 | 4 | 36.4 | 0.44 | 0.950 | 1.00 |
| Pyridines and derivatives | 27 | 21 | 77.8 | 1.19 | 0.457 | 0.984 | 14 | 51.9 | 0.83 | 0.753 | 1.00 |
| Fatty acids and conjugates | 32 | 24 | 75.0 | 1.02 | 0.579 | 1.00 | 18 | 56.2 | 0.99 | 0.583 | 0.993 |
| 1,2-diacylglycerols | 17 | 10 | 58.8 | 0.48 | 0.957 | 1.00 | 8 | 47.1 | 0.68 | 0.848 | 1.00 |
| 1,3-diacylglycerols | 31 | 17 | 54.8 | 0.41 | 0.996 | 1.00 | 13 | 41.9 | 0.55 | 0.965 | 1.00 |
| 1-acyl-sn-glycero-3-phosphoethanolamines | 26 | 20 | 76.9 | 1.13 | 0.502 | 1.00 | 12 | 46.2 | 0.66 | 0.896 | 1.00 |
| 2-acyl-sn-glycero-3-phosphocholines | 11 | 8 | 72.7 | 0.90 | 0.706 | 1.00 | 4 | 36.4 | 0.44 | 0.950 | 1.00 |
| Alpha amino acids | 33 | 15 | 45.5 | 0.28 | 1.00 | 1.00 | 10 | 30.3 | 0.33 | 0.999 | 1.00 |
| Amino acids, peptides, and analogues | 84 | 55 | 65.5 | 0.63 | 0.979 | 1.00 | 44 | 52.4 | 0.84 | 0.809 | 1.00 |
| Azolidines | 12 | 9 | 75.0 | 1.02 | 0.640 | 1.00 | 4 | 33.3 | 0.38 | 0.972 | 1.00 |
| Benzene and substituted derivatives | 12 | 7 | 58.3 | 0.47 | 0.943 | 1.00 | 6 | 50.0 | 0.77 | 0.773 | 1.00 |
| Benzodioxoles | 14 | 10 | 71.4 | 0.85 | 0.733 | 1.00 | 7 | 50.0 | 0.77 | 0.777 | 1.00 |
| Benzoic acids and derivatives | 16 | 12 | 75.0 | 1.02 | 0.620 | 1.00 | 7 | 43.8 | 0.60 | 0.899 | 1.00 |
| Coumarins and derivatives | 20 | 14 | 70.0 | 0.79 | 0.777 | 1.00 | 7 | 35.0 | 0.41 | 0.985 | 1.00 |
| Dipeptides | 160 | 91 | 56.9 | 0.43 | 1.00 | 1.00 | 56 | 35.0 | 0.40 | 1.00 | 1.00 |
| Diphenylmethanes | 10 | 6 | 60.0 | 0.51 | 0.919 | 1.00 | 5 | 50.0 | 0.77 | 0.769 | 1.00 |
| Fatty amides | 15 | 7 | 46.7 | 0.29 | 0.996 | 1.00 | 6 | 40.0 | 0.51 | 0.939 | 1.00 |
| Gamma amino acids and derivatives | 10 | 7 | 70.0 | 0.79 | 0.769 | 1.00 | 3 | 30.0 | 0.33 | 0.978 | 1.00 |
| Gluco/mineralocorticoids, progestogins and derivatives | 14 | 7 | 50.0 | 0.34 | 0.989 | 1.00 | 4 | 28.6 | 0.31 | 0.992 | 1.00 |
| Glutamine and derivatives | 15 | 4 | 26.7 | 0.12 | 1.00 | 1.00 | 3 | 20.0 | 0.19 | 0.999 | 1.00 |
| Hydroxycinnamic acids and derivatives | 11 | 3 | 27.3 | 0.13 | 1.00 | 1.00 | 4 | 36.4 | 0.44 | 0.950 | 1.00 |
| L-alpha-amino acids | 19 | 11 | 57.9 | 0.46 | 0.969 | 1.00 | 6 | 31.6 | 0.35 | 0.992 | 1.00 |
| Lineolic acids and derivatives | 65 | 31 | 47.7 | 0.30 | 1.00 | 1.00 | 23 | 35.4 | 0.42 | 1.00 | 1.00 |
| Monoterpenoids | 15 | 11 | 73.3 | 0.93 | 0.677 | 1.00 | 5 | 33.3 | 0.38 | 0.981 | 1.00 |
| N-acyl-alpha amino acids | 34 | 22 | 64.7 | 0.62 | 0.935 | 1.00 | 19 | 55.9 | 0.98 | 0.598 | 1.00 |
| Peptides | 15 | 10 | 66.7 | 0.68 | 0.846 | 1.00 | 7 | 46.7 | 0.67 | 0.848 | 1.00 |
| Peptidomimetics | 15 | 6 | 40.0 | 0.22 | 0.999 | 1.00 | 6 | 40.0 | 0.51 | 0.939 | 1.00 |
| Thiazoles | 41 | 18 | 43.9 | 0.26 | 1.00 | 1.00 | 14 | 34.1 | 0.40 | 0.999 | 1.00 |
| Triacylglycerols | 60 | 34 | 56.7 | 0.44 | 0.999 | 1.00 | 9 | 15.0 | 0.13 | 1.00 | 1.00 |
| Tricarboxylic acids and derivatives | 16 | 11 | 68.8 | 0.74 | 0.803 | 1.00 | 4 | 25.0 | 0.26 | 0.998 | 1.00 |
| Vitamin D and derivatives | 16 | 12 | 75.0 | 1.02 | 0.620 | 1.00 | 9 | 56.2 | 0.99 | 0.610 | 1.00 |

*Notes:* Classes are ordered by the smaller of their two *q* values. In class, number of the 3,820 annotated metabolites assigned to the class. Assoc. (n) and Assoc. (%), number and percentage of the class significantly correlated with the shared-factor score (Spearman correlation, Benjamini–Hochberg FDR < 0.05, controlled jointly across microbial and metabolomic features and both factors); the percentage in each column heading is the value expected for a class under no enrichment. Odds ratio, (*a* × *d*)/(*b* × *c*) for the 2 × 2 table of class membership by factor association, where *a* and *b* are the associated and non-associated metabolites in the class and *c* and *d* those outside it; values > 1 indicate over-representation. *P*, one-sided hypergeometric test for over-representation. *q*, Benjamini–Hochberg-adjusted *P* across the 97 classes, computed separately for each factor; *q* < 0.05 in bold. <sup>a</sup>Odds ratio computed with the Haldane–Anscombe correction (0.5 added to each cell) because one cell of the 2 × 2 table was zero: Ergosterols and derivatives, Naphthalenes, Phenylbenzodioxanes (SF<sub>1</sub>); Carboximidic acids and derivatives (SF<sub>2</sub>). FDR, false discovery rate; SF, shared factor.

